# Benchmarking algorithms for gene regulatory network inference from single-cell transcriptomic data

**DOI:** 10.1101/642926

**Authors:** Aditya Pratapa, Amogh P. Jalihal, Jeffrey N. Law, Aditya Bharadwaj, T. M. Murali

## Abstract

We present a comprehensive evaluation of state-of-the-art algorithms for inferring gene regulatory networks (GRNs) from single-cell gene expression data. We develop a systematic framework called BEELINE for this purpose. We use synthetic networks with predictable cellular trajectories as well as curated Boolean models to serve as the ground truth for evaluating the accuracy of GRN inference algorithms. We develop a strategy to simulate single-cell gene expression data from these two types of networks that avoids the pitfalls of previously-used methods. We selected 12 representative GRN inference algorithms. We found that the accuracy of these methods (measured in terms of AUROC and AUPRC) was moderate, by and large, although the methods were better in recovering interactions in the synthetic networks than the Boolean models. Techniques that did not require pseudotime-ordered cells were more accurate, in general. The observation that the endpoints of many false positive edges were connected by paths of length two in the Boolean models suggested that indirect effects may be predominant in the outputs of the algorithms we tested. The predicted networks were considerably inconsistent with each other, indicating that combining GRN inference algorithms using ensembles is likely to be challenging. Based on the results, we present some recommendations to users of GRN inference algorithms, including suggestions on how to create simulated gene expression datasets for testing them. BEELINE, which is available at http://github.com/murali-group/BEELINE under an open-source license, will aid in the future development of GRN inference algorithms for single-cell transcriptomic data.

---

It is now possible to measure the transcriptional states of individual cells in a sample due to advances in single-cell RNA-sequencing technology. Although these measurements are noisy, they permit several new applications. One prominent direction is clustering of cells based on shared expression patterns leading to the discovery of new cell types [1, 2]. A central question that arises now is whether we can discover the gene regulatory networks (GRNs) that control cellular differentiation into diverse cell types and drive transitions from one cell type to another. Single-cell expression data are especially promising to address this challenge because, unlike bulk transcriptomic data, they do not obscure biological signals by averaging over all the cells in a sample.

Not surprisingly, over a dozen methods have been developed or used to infer GRNs from single-cell data [3–15]. Some of these methods were originally developed for bulk transcriptional data but have since been applied to single-cell data [3, 4, 13, 16]. Other publications have presented approaches to reverse engineer Boolean networks from single cell transcriptomic data [11, 17–19]. The criteria for evaluation and comparison of algorithms often change from one paper to another. Moreover, there are no widely-accepted ground truth datasets for assessing the accuracy of these methods. As a result, an experimentalist seeking to analyze a new dataset faces a daunting task in selecting an appropriate inference method. Which method is appropriate for my dataset? How should its parameters be tuned? What might the running time be? How can I test the accuracy of the resulting network? Given the rapid developments in this field, it is critical as well as timely to design a comprehensive evaluation framework to assess the accuracy, robustness, and efficiency of GRN inference methods based on clearly-defined benchmark datasets.

There are three challenges that arise when we evaluate GRN inference algorithms for single-cell RNA-seq data. First, the “ground truth”, i.e. the network of regulatory interactions governing the dynamics of genes of interest, is usually unknown. Consequently, it is a common practice to create artificial graphs or extract subnetworks from large-scale transcriptional networks. This procedure raises the second challenge: there is no accepted strategy to accurately simulate single-cell gene expression data from these networks. Methods such as GeneNetWeaver [20] are widely used to synthesize bulk transcriptomic data from GRNs. GeneNetWeaver has also been applied in single-cell analysis [7, 9, 10, 18, 21] but has limitations, as we demonstrate. Third, many algorithms need temporal ordering of cells as part of the input. It is quite often the case that the single cell experiment used to generate the gene expression data does not provide this information. While pseudotime is the most popular surrogate for experimental time, there are nearly a hundred techniques available for estimating it [22] and their limitations may propagate to inaccuracies in GRN reconstruction. We use multiple and distinct strategies to address these challenges.

To address the first challenge, we start with two sets of networks that serve as the ground truth for GRN inference. The first group includes six “toy” networks with specific topologies that give rise to different cellular trajectories with predictable qualitative properties, e.g., linear, cyclic, bifurcating, and converging [22]. The nodes in these synthetic networks do not correspond to real genes. For the second set of networks, we curate four published Boolean models that explore gene regulatory interactions underlying various developmental and tissue differentiation processes. Mutual inhibition between a pair of genes is a key characteristic of each of these models; this type of relationship is important in creating branching gene expression trajectories. The regulatory networks underlying the Boolean models serve as the ground truth during evaluation.

To deal with the second challenge, we develop a method called BoolODE that converts a Boolean model into a system of ordinary differential equations (ODEs). We expand the Boolean function combining the regulators of each gene into a truth table in order to infer the form of the corresponding ODE representation. This approach provides a reliable method to capture the logical relationships among the regulators precisely in the components of the ODE. We add noise terms to make these equations stochastic [22, 23]. We demonstrate that the gene expression datasets arising from our simulations capture exactly as many distinct trajectories as the number of final states exhibited by the synthetic network or the Boolean model.

To tackle the third challenge, we use two ideas. For datasets from synthetic networks, we use the simulation time as input to the GRN algorithms so that we can isolate them from the limitations of pseudotime calculation. In the case of datasets from curated Boolean models, we check that the pseudotime values we compute are well correlated with the simulation times.

We combine all these strategies in BEELINE, a comprehensive framework for evaluating methods that infer gene regulatory networks from single-cell gene expression data (Figure 1). BEELINE incorporates 12 diverse GRN inference algorithms. It provides an easy-to-use and uniform interface to the implementation of each inference method in the form of a Docker image. This flexibility makes it very easy to include a new GRN inference technique in the overall framework. BEELINE implements several methods for estimating and comparing the accuracy, stability, and efficiency of these algorithms.Thus, BEELINE facilitates reproducible, rigorous, and extensible evaluations of GRN inference methods for single-cell gene expression data.

**Figure 1:**
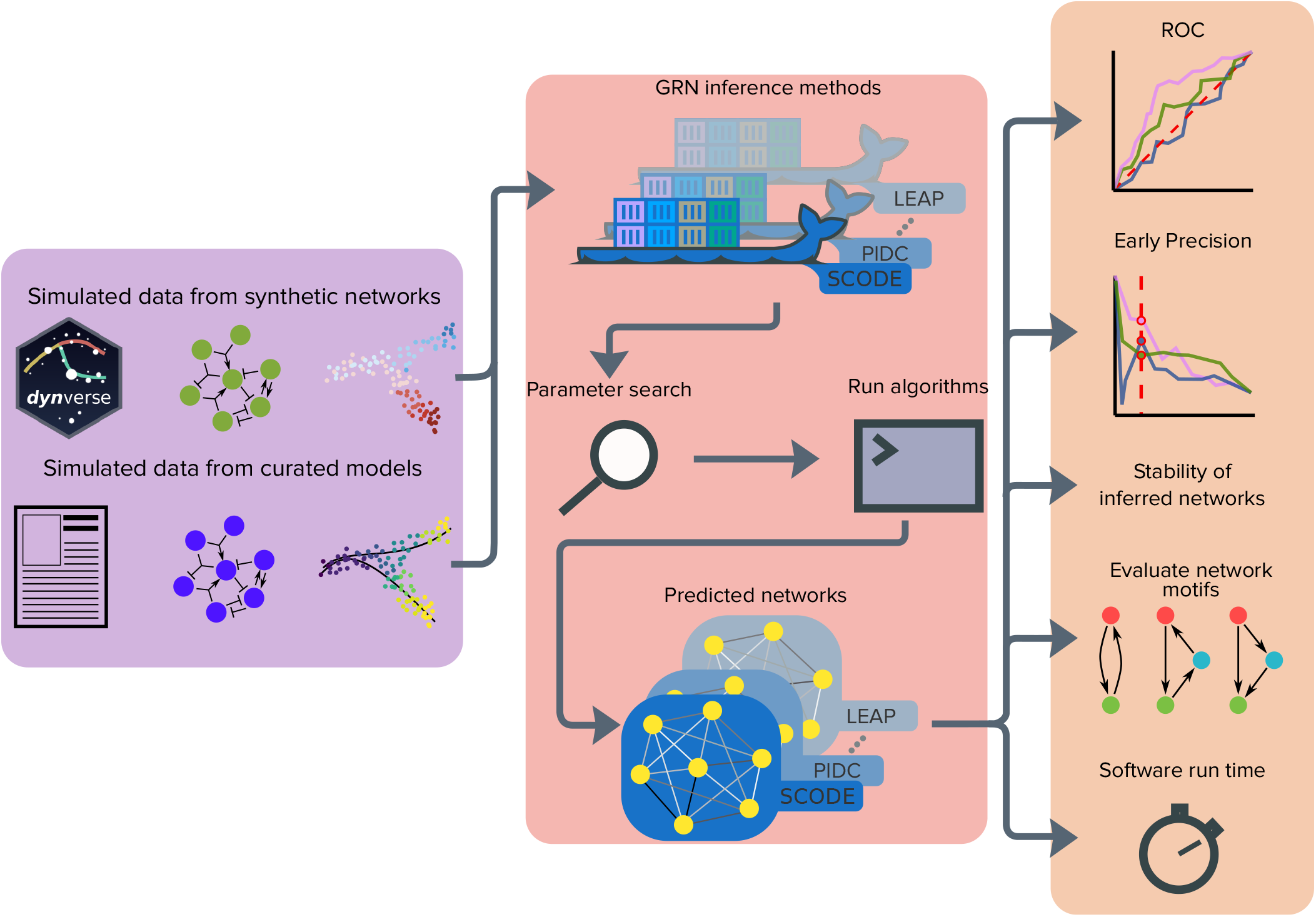
An overview of the BEELINE evaluation framework. We apply GRN inference algorithms to two types of data: datasets from synthetic networks and datasets from curated models from the literature. We process each dataset through a uniform pipeline: pre-processing, Docker containers for 12 GRN inference algorithms, parameter estimation, post-processing, and evaluation. We compare algorithms based on accuracy (ROC and precision-recall curves), stability of results (correlations across simulations, similarities and differences across algorithms), visualization, analysis of network motifs, and running time.

## Results

### Overview of Algorithms

We surveyed the literature for papers that either published a new algorithm on reverse engineering gene regulatory networks or used an existing algorithm. We included three preprints deposited in bioRxiv. We required that each paper applied the algorithm to at least one experimental single-cell transcriptional dataset. We did not place any constraint on the type of experimental platform used to collect the data analyzed. We ruled out methods that output a network without assigning a weight or rank to the interactions or whose output changed considerably from one run to another. We did not consider methods that required additional datasets or supervision to compute GRNs or whose goal was to discover cell-type specific networks. We checked if the software was available for download and was easy to install.

We selected 12 methods using these criteria. We briefly describe each algorithm in “Regulatory Network Inference Algorithms”. Most algorithms developed explicitly for single-cell transcriptomic data required the cells to be ordered by pseudotime in the input, with PIDC [7] being an exception. These methods ideally require datasets corresponding to linear trajectories; some techniques recommend that data with branched trajectories be split into multiple linear ones before input [12, 15]. In contrast, methods that had originally been developed for bulk transcriptional data did not impose this requirement [3, 4]. Almost all the methods we included output directed networks with exceptions being PPCOR and PIDC [4, 7]. Only five methods output signed networks, i.e., they indicated whether each interaction was activating or inhibitory [4, 6, 9–11]. A number of methods inferred each pairwise interaction independently of the others, sometimes conditioned on the other genes [4, 5, 7, 12]. Several other methods computed all the regulators of a gene simultaneously but solved the problem independently for each gene [3, 6, 9, 11, 13–15]. The first few columns of Figure 3 summarize these properties.

We use BEELINE to evaluate the 12 inference algorithms on 300 simulated datasets across six synthetic networks and four curated Boolean models. We first compare the algorithms for datasets from synthetic networks before presenting results for datasets from curated models.

### Datasets from Synthetic Networks

Our motivations for generating *in silico* single-cell gene expression datasets from synthetic networks were two-fold. First, we wanted to use a known GRN that could serve as the ground truth for the evaluation of the accuracy of the networks proposed by each method. Second, since several GRN inference methods require some kind of time ordering of the cells in the input, we desired to create datasets that were isolated from any limitations of pseudotime inference algorithms.

To this end, we started with six synthetic networks (Figure 2(a)). Saelens *et al.*[22] have argued that simulating these networks should produce a variety of different trajectories that are commonly observed in differentiating and developing cells, namely, Linear, Linear Long, Cycle, Bifurcating, Bifurcating Converging, and Trifurcating.

**Figure 2:**
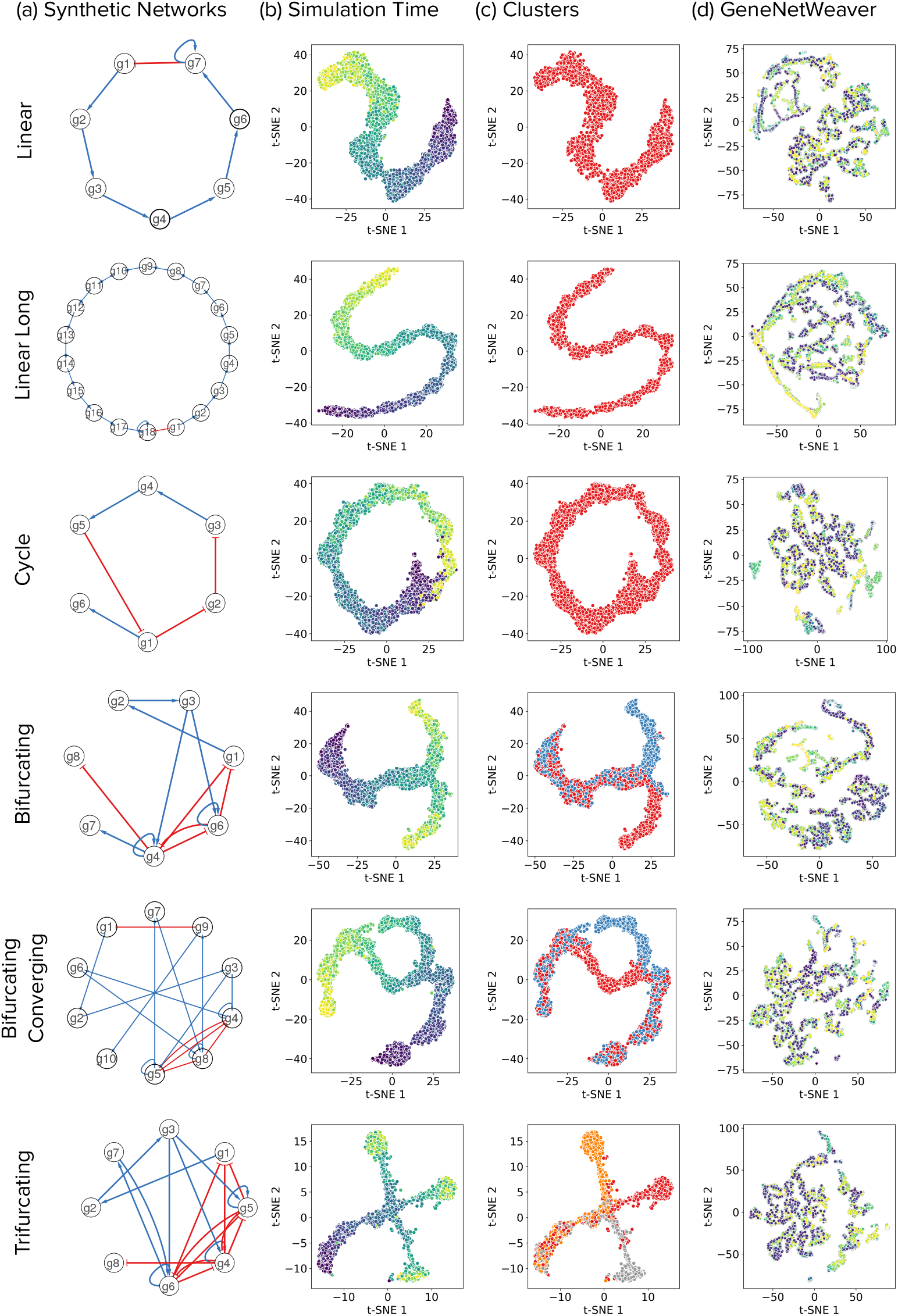
A comparison of datasets from synthetic networks simulated using BoolODE and GeneNetWeaver. Each row corresponds to the network indicated by the label on the left. (a) The network itself, with red edges representing inhibition and blue edges representing activation. (b) A 2D t-SNE visualization of the BoolODE output. The colour of each point indicates the simulation time: blue for earlier, green for intermediate, and yellow for later times. (c) Each colour corresponds to a different subset of cells obtained by using *k*-means clustering of simulations, with *k* set to the number of expected steady states. (d) A 2-D t-SNE visualization of the GeneNetWeaver output.

#### Generation of simulated datasets

We considered how to simulate these networks to create in *silico* single-cell gene expression datasets. Several recent studies on GRN inference from such data [7, 9, 10, 18, 21] have used GeneNetWeaver [24], a method originally developed for generating time courses of bulk-RNA datasets from a given GRN. Accordingly, we simulated the six synthetic networks using GeneNetWeaver via the procedure outlined in Chan *et al.* [7] (see Supplementary Section S1 for details). However, we observed no discernible trajectories when we visualized two-dimensional projections of these data (Figure 2(d)).

We therefore used our BoolODE approach to simulate the six synthetic networks; we describe BoolODE in Supplementary Section S1. We obtained 30 different datasets by varying the number of cells (10 each with 500, 2000 and 5000 cells) for each of the networks using the procedure outlined in “Creating Datasets from Synthetic Networks”. We visually inspected the two-dimensional representation of these datasets computed by t-SNE and confirmed that the simulation of each synthetic network yielded the expected trajectory (Figure 2(b)). Next, we clustered the simulations in each dataset using *k*-means clustering with *k* set to the expected number of trajectories (see “Creating Datasets from Curated Models” for details). Each colour in the t-SNE plots Figure 2(c) corresponds to a distinct cluster. Visually comparing the corresponding plots in Figures 2(b) and (c), we confirmed that each cluster indeed included cells from a distinct trajectory. These results reassured us that BoolODE was successful in correctly simulating the network models.

We executed each of the 12 algorithms using each of the 30 simulated single-cell datasets and the corresponding simulation times for the six synthetic networks as input. For those GRN inference methods that required time information for each cell, we provided the simulation time at which we sampled each cell. For the Bifurcating, Bifurcating Converging and Trifurcating networks, we ran the algorithms that need time information on each trajectory individually and combined the outputs using the procedure outlined in “Evaluation Pipeline”.

#### Evaluation of accuracy of inferred GRNs

We used the following approach to evaluate the GRNs computed by each algorithm. We set each synthetic network as the ground truth. We compared the GRN for each of the 30 simulated datasets for that network against the ground truth. We plotted ROC and precision-recall curves and measured the areas under these curves. We computed the median AUROC and AUPRC values for each synthetic network across the 30 different datasets. We also sought to estimate the stability of the results of each algorithm as we varied the samples and the number of cells. Therefore, we determined the “top-*k*” network for each GRN algorithm, with *k* set to the number of edges in each ground truth network. We computed the Jaccard index of these sets of edges for all 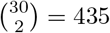 pairs of datasets and recorded the median values. See “Evaluation Pipeline” for details.

Figure 3 summarizes our results while Supplementary Figures S1 and S2 show the box plots of AUROC and AUPRC values. In Figure 3, we ordered the algorithms by the median of the per-network median AUROC values. We observed that 10 out of 12 methods had a median AUROC greater than 0.75 on the Linear network, with SINCERITIES and GRISLI achieving a median AUROC as high as 0.90. All 12 methods had a median AUROC greater than 0.65 in the Linear Long Network, with four methods achieving a median AUROC greater than 0.90. SINCERITIES had the highest median AUROC in three out of the six synthetic networks, namely, Linear (0.90) and Bifurcating (0.82), and tied with SCINGE for Cycle (0.93). GRNVBEM had the highest median AUROC (0.96) in the Linear Long network, SCINGE (0.83) in the Bifurcating Converging and LEAP (0.73) in the Trifurcating network. While most methods achieved a higher-than-random AUROC (0.5) in general, we noted that GRNVBEM for the Cycle network and SCODE for the Trifurcating network have a median AUROC less than 0.5. However, we did not see a similar ordering in terms of AUPRC values. The AUPRC values rarely reached 0.75 with a large fraction being 0.5 or less. Examining Supplementary Figures S1 and S2, we did not observe any clear improvement in AUROC or AUPRC as we the increased number of cells in the datasets.

**Figure 3:**
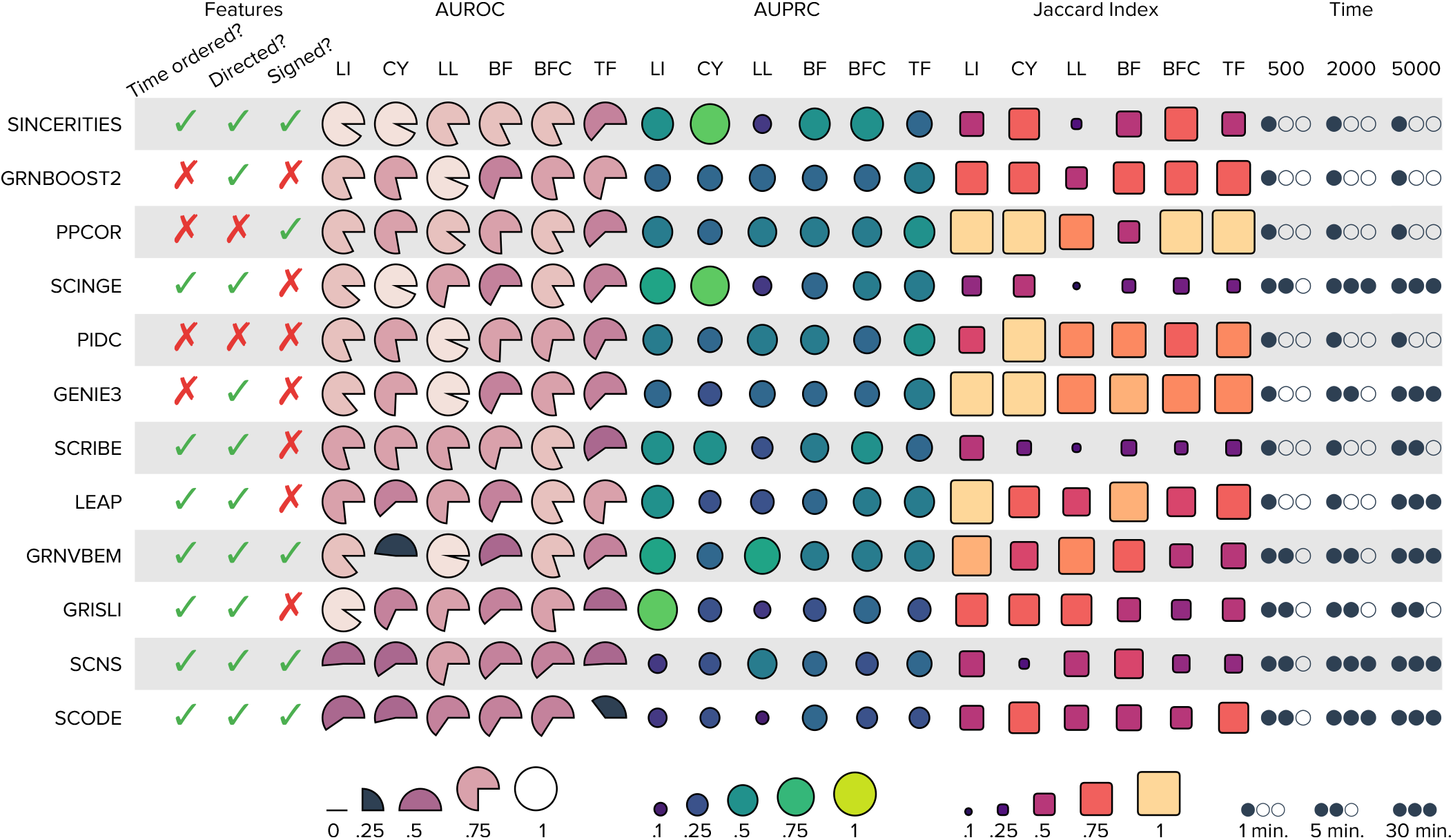
Summary of results for datasets from synthetic networks. Each row corresponds to one of the algorithms included in our evaluation. The first three columns record properties of each algorithm: whether it requires time-ordered cells, whether it outputs a directed network, and whether each edge in the output has a sign. The next three sets of six columns displays the median area under the ROC curve, the median area under the precision-recall curve, and the median Jaccard index, respectively, for each of the six simulated datasets. The area of a shape is proportional to the corresponding value between 0 and 1. A blackened wedge represents median AUROC values less than 0.50. The final set of three columns shows the mean running time. We computed the medians and means over 30 simulated datasets for each synthetic network. Abbreviations: LI: Linear, CY: Cycle, LL: Linear Long, BF: Bifurcating, BFC: Bifurcating Converging, TF: Trifurcating.

#### Stability analysis

While SINCERITIES had the highest median AUROC scores across a majority of the networks, the top-*k* edges predicted by SINCERITIES were relative less stable with a median Jaccard index as low as 0.24 (for the Long Linear network, which is the most sparse) and indices around 0.5 for three other synthetic networks. Among the top-three methods with the highest median-of-median AUROC values, PPCOR achieved the highest median Jaccard index of 1.0 for four out of six synthetic networks. This result suggests that partial correlation may be robust to variations in sampling and dataset size. The stability of GRNBoost2 was intermediate between PPCOR and SINCERITIES, with the Jaccard index being close to 0.75 for as many as five networks. We noted that Jaccard index was relatively low for SCINGE and SCRIBE with the maximum value across the six networks being 0.55 for SCINGE and 0.44 for SCRIBE.

#### Running time

We also recorded the running times of these algorithms (mean time shown in the last three columns of Figure 3). We observed that seven of the 12 GRN algorithms were fairly fast in that they took under a minute on average on datasets with 500 cells, while five others ran in under five minutes on average. We also observed that the computation time was under a minute irrespective of the number of cells for four methods (SINCERITIES, GRNBoost2, PPCOR, and PIDC). However, in general, the running time of GRN inference methods increased with the dataset size, with six out of 12 methods taking around 30 minutes to an hour on average when provided with a dataset of 5,000 cells.

### Datasets from Curated Models

Synthetic networks or dense sub-networks of large-scale gene regulatory networks are primarily used to generate simulated datasets of single-cell gene expression [7, 9, 10, 18, 21]. However, these networks may not capture the complex regulation in any specific developmental process. To avoid these pitfalls, we curated the literature for published Boolean models that explore gene regulatory interactions in various developmental processes. A benefit of using these Boolean models is that they could serve as the ground-truth regulatory networks to evaluate the performance of proposed GRNs. We collected four Boolean models for Mammalian Cortical Area Development (mCAD) [25], Ventral Spinal Cord Development (VSC) [26], Hematopoietic Stem Cell Differentiation (HSC) [27], and Gonadal Sex Determination (GSD) [28]. Figure 4(a) displays these networks.

**Figure 4:**
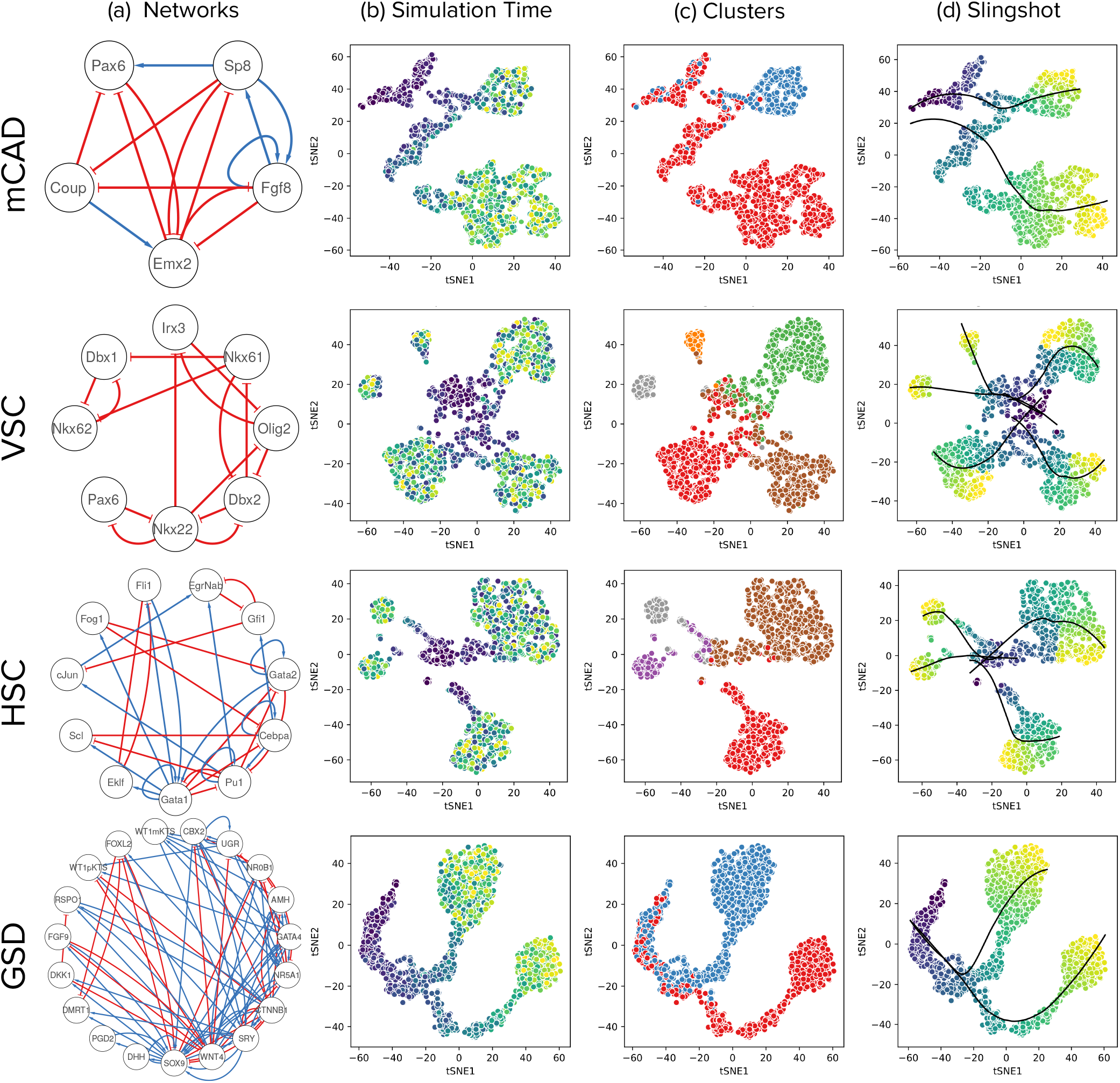
t-SNE visualization reveals trajectories leading to states that correspond to those of the Boolean model. Each row in the figure corresponds to a Boolean model, indicated on the left. (a) Network diagrams of the models. (b) The t-SNE visualizations of cells sampled from BoolODE output. The color of each point indicates the corresponding simulation time (blue for early, green for middle, and yellow for later times). (c) Each colour corresponds to a different subset of cells obtained by using *k*-means clustering of simulations, with *k* set to the number of steady states reported in the relevant publication. (d) Slingshot pseudotime output. The principal curves obtained using Slingshot are shown in black.

#### Correspondence of BoolODE simulations and steady states of Boolean models

In the case of datasets from synthetic networks, we were able to visualize clear correspondences between the simulated data and expected trajectories from the networks. In the case of Boolean models, we sought to determine if the simulated datasets could capture steady state properties of the Boolean models, as reported in the corresponding publications. We performed this analysis in two stages.

i. Figure 4(b) shows the two-dimensional t-SNE visualizations colored with simulation time for one of 10 datasets created by BoolODE for each Boolean model. Examining these plots visually, we observed continuous, discernible trajectories that ran along the entire simulation timeline, as seen by the colours of the cells changing gradually from blue to green to yellow. Next, we carried out *k*-means clustering of simulations with *k* set to the number of steady states of the model, as mentioned in the corresponding publication. Figure 4(c) displays these results, with a different color for each cluster (two for mCAD, five for VSC, four for HSC, and two for GSD). We visually corresponded the plots in Figures 4(b) and (c) to confirm that each cluster contained cells spanning the entire length of the simulation. After running Slingshot [29], we visualized the principal curves computed by this method in Figure 4(d); each such curve computed is a smooth representation of the trajectory. The correlation between the pseudotime and the simulation time was high (0.84 for mCAD, 0.61 for VSC, 0.8 for HSC, and 0.93 for GSD). Therefore, we gave the pseudotime as input to the GRN algorithms.
ii. Each publication describing a Boolean model specifies a unique gene expression pattern that characterizes each steady state of that model. We asked if we our simulated data matched these patterns. To this end, we colored each cell (a point on the 2D t-SNE plot) with the simulated expression value of each gene in that cell, with darker colors corresponding to larger values (Supplementary Figures S3 to S6). We explain the procedure we followed using an example. The mCAD model has two steady states: Anterior (characterized by up-regulation of Pax6, Fgf8, and Sp8) and Posterior (up-regulation of Coup and Emx2). In Supplementary Figure S3, we observed that Pax6, Fgf8, and Sp8 had high simulated expression values in a group of cells at the top right of the t-SNE plot whereas Coup and Emx2 had large values in the group in the bottom left. We concluded that these two cell groups corresponded to the Anterior and Posterior steady states of the Boolean model, respectively. We succeeded in finding similar matches for each of the other models (Supplementary Figures S4 to S6).

#### Generation of simulated datasets

Encouraged by these results, we used BoolODE to create 10 different datasets with 2,000 cells for each model. We computed pseudotime using Slingshot [29] and provide these values to the GRN algorithms, so as to mimic a real single-cell gene expression data processing pipeline. For each dataset, we also generated a version that included varying levels of dropouts (*q*) [7]. We obtained 30 datasets for each of the four models, 10 datasets with no dropout, 10 datasets with dropouts at *q* = 20, and 10 datasets with dropouts at *q* = 50. We provide details of dataset generation in “Pre-processing Datasets from Curated Models”. We then ran each of the 12 GRN inference algorithms and analyzed each of the 30 reconstructions proposed by these methods for the four curated models.

#### Evaluation of accuracy of inferred GRNs

We compared the proposed GRN reconstructions to the ground truth network for each the Boolean models using a similar approach as with the datasets from synthetic networks. In addition to computing AUROC and AUPRC, we computed the early precision of the top-*k* edges as well as “signed” versions of early precision separately for activating and inhibitory edges in the ground truth network. We also computed the early precision ratio (EPR) by dividing the early precision score with the the expected precision of a random predictor, which is the edge density of the ground truth network. An EPR value of one means that a method has an early precision score equal to that of a random predictor. We recorded the medians of these values across the 10 datasets for each Boolean model. Figure 5 summarizes our findings. As with datasets from synthetic networks, we ordered the algorithms by the median of the per-model median AUROC values. We present boxplots of the 10 AUROC values for each algorithm and each dataset in Supplementary Figure S7.

**Figure 5:**
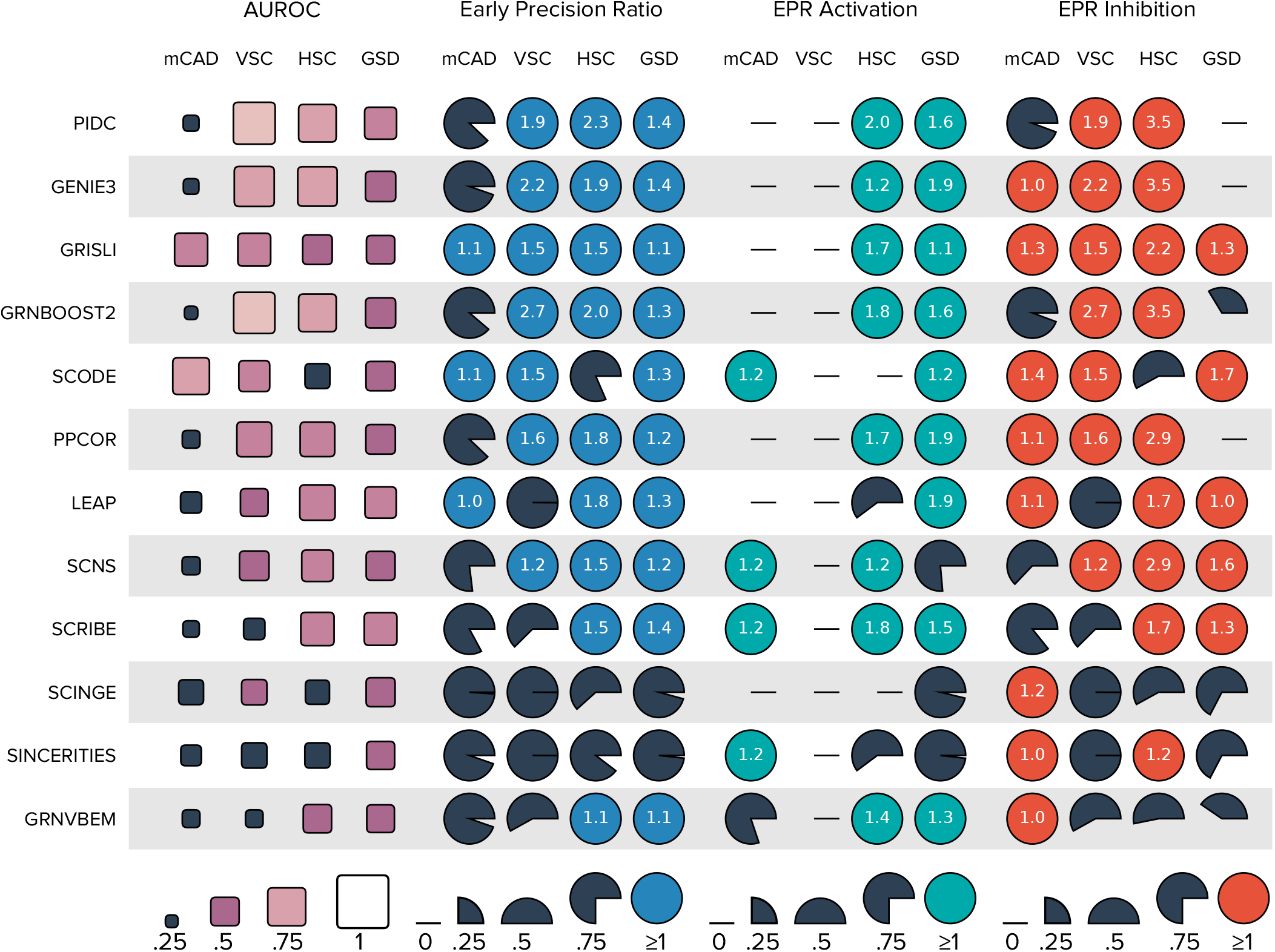
Summary of results for datasets from curated models. Each row corresponds to one of the algorithms included in our evaluation. The four sets of four columns each display the median AUROC, median early precision ratio (EPR), median EPR for activating edges, and median EPR for inhibitory edges. When the EPR is at least one, it appears in the circle. A blackened square or wedge represents values that are less than that of a random predictor.

#### AUROC results

We focus on the 10 datasets without dropouts (first four columns of Figure 5). We observed that only two (GRISLI and SCODE) out of the 12 methods had a median AUROC greater than 0.50 for the mCAD model, which has a network density 0.65 (excluding self-loops). The high density of the network proved quite challenging for almost every method. For the VSC model, which only has inhibitory edges, three methods (PIDC, GRNBoost2, and GENIE3) had an AUROC greater than 0.75. The same three methods had AUPRC close to 0.75 for the HSC model. SCRIBE (0.64), PIDC (0.53), GENIE3 (0.59), and GRNBoost2 (0.59) had the four highest AUROC values for the GSD model. Overall, we observed that GENIE3, GRNBoost2, and PIDC had among the highest median AUROCs for three out of the four models. Surprisingly, SINCERITIES, which was the best algorithm according to the AUROC values for the datasets from synthetic networks, had a close to random median AUROC for three out of the four curated models. For datasets with dropouts, we did not see any clear trends in terms of smaller or larger AUROC values compared to the dropout-free results (Supplementary Figure S7).

#### Early precision results

Next, we studied the early precision and EPR values of the top-*k* predictions for each of the four models. Columns five through eight of Figure 5 show the median early precision ratios (EPRs) for each method. We observed that 11 algorithms had a median early precision worse than or close to a random predictor in at least one of the four models (black wedges in Figure 5). In the 29 cases when the EPR was at least one, it was 1.5 or larger only 14 times. With a network density of 0.65, the mCAD model had the smallest number of algorithms with larger early precision than a random predictor. For this model, only three algorithms (GRISLI, LEAP, SCODE) had an early precision greater than this network density. For the VSC model, which has a network density of 0.27, five methods had a worse-than-random early precision. The HSC and GSD have comparable network densities as VSC (0.24 and 0.26, respectively), but fewer than four methods had an early precision less than the baseline values for both these models. Thus, network density alone was not a good predictor of early precision values. For datasets with dropouts, we did not see any clear trends in terms of smaller or larger early precision values compared to the dropout-free results (Supplementary Figure S8).

#### Results on activating and inhibitory edges

Next, we investigated if the GRN inference methods were better at recovering activating edges or inhibitory edges. We computed the early precision of signed edges using the procedure outlined in “Evaluation Pipeline”. Since the VSC model only contains inhibitory edges, the EPR for activating edges does not apply to it. The mCAD model was an outlier for activating edges, since it has only five such interactions. Seven methods had very poor EPR values. SCODE, SCNS, SCRIBE, and SINCERITIES had slightly better-than-random scores. In contrast, for inhibitory interactions, eight methods performed slightly better than a random predictor. Overall, the GRN inference algorithms we evaluated tend to perform poorly when it comes to recovering the true edges within the top-*k* predictions.

#### Network motifs

We investigated if certain network motifs were over-represented in the top-*k* networks. Specifically, we counted the number of three-node feedback and feedforward loops and (two-node) mutual interactions in the top-*k* networks from each GRN algorithm and reported their ratios with their respective values in the ground truth network. We report the medians of these values (over the 10 datasets we simulated for each model) in Figure 6(a). For the mCAD model, we observed that all GRN inference methods yielded close to expected or lower than expected number of each network motif compared to the ground truth network. This trend is likely due to the fact that the mCAD network is fairly dense. For the other networks, we noted an overall decrease in feedback loops and mutual interactions but an increase in feedforward loops. In the case of feedback loops, we noted that SCRIBE was the only method that predicted roughly as many or more feedback loops than in the ground truth network for each of the four Boolean models.

**Figure 6:**
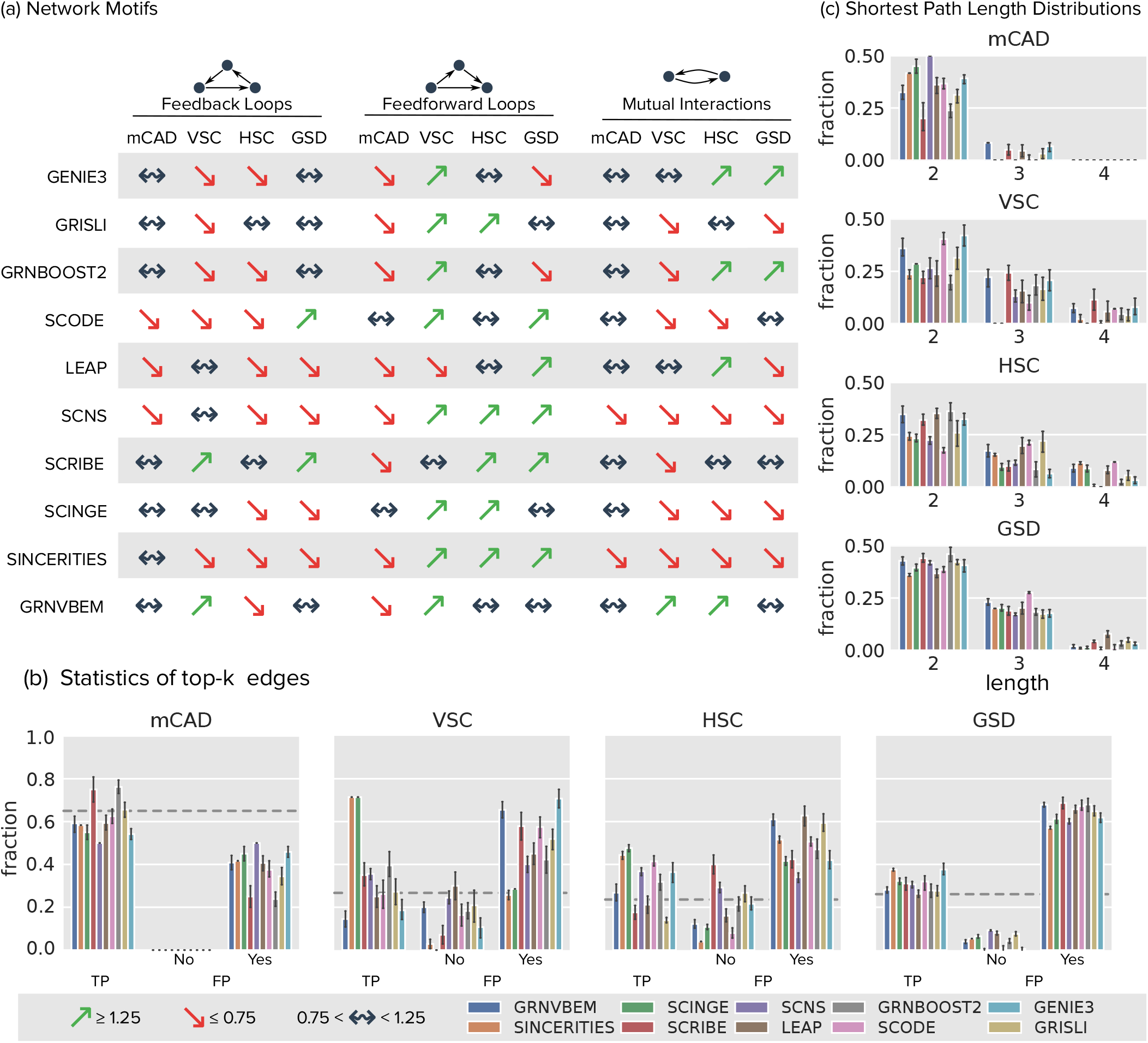
(a) Analysis of network motifs in the predicted top-*k* networks using datasets from curated models. We considered three motifs, namely, feedback loops, feedforward loops and mutual interactions. We counted the number of occurrences of each motif type in each predicted top-*k* network, as well as in the ground truth network, and we report the ratio of these counts. Green upward pointing arrows represent ratios greater than 1.25, red downward pointing arrows represent ratios less than 0.75, and the black arrows represent ratios between 0.75 and 1.25. (b) Statistics on the edges in the top-*k* network for each method divided into three parts: fraction of top-*k* edges that are true positives (TP), fraction of top-*k* edges that are false positives and whose endpoints are not connected by any path in the ground truth network (FP, No), and the fraction of top-*k* edges that are false positives and whose endpoints are connected by a path (of length two or more) in the ground truth network (FP, Yes). Each colored bar corresponds to one algorithm. The height of a bar represents the mean and the error bar the 95% confidence interval over the 10 datasets. The gray line indicates the density of the network. (c) For every false positive edge in each predicted top-*k* network, we computed the length of the shortest paths in the ground truth network that connected the endpoints of this edge. Bar plots show the fractions of top-*k* edges that are false positive edges and whose endpoints have paths of lengths 2, 3, and 4 in the ground truth network in the top-*k* networks, colored by the method.

#### Explaining false positives in top-*k* networks

We reasoned that the poor value of early precision for a GRN inference algorithm could be due to its tendency to predict “indirect” interactions, i.e., if, in the Boolean model, gene *a* regulates gene *b*, which in turn regulates gene *c*, a method may also predict an interaction between *a* and *c*. This type of configuration is a feedforward loop, which we have seen to be in excess over the ground truth network for several algorithms (Figure 6). To confirm the possibility of indirect interactions, we considered each false positive edge (*u, v*) in a top-*k* network and computed the length of the shortest path between *u* and *v* in the corresponding ground truth network. Figures 6(b) and (c) summarize the results.

We discuss the results for the VSC, HSC, and GSD models first. We observed that the endpoints of about 10% to 20% of false positive edges were not even connected in the ground truth networks by any path (bars labeled “No” in Figure 6(b)). Virtually all the remaining false positive edges corresponded to paths of lengths between two and four in the ground truth network (Figure 6(c)); this figure does not show a miniscule fraction of false positive edges that corresponded to paths of length five in the GSD network. What was most striking was that the largest proportion of these false positive edges were connected by paths of length two, i.e., if added to the ground truth network, they would indeed form feed-forward loops. This result lends strong credence to our hypothesis that GRN inference algorithms are predicting indirect interactions.

In the case of the mCAD model, a very large fraction of false positive edges corresponded to paths of length two in the corresponding ground truth network. We reasoned that the density of the mCAD network was the cause for this statistic despite the lower prevalence of feedforward loops in the predicted networks for the mCAD model.

#### Comparing similarities of algorithm outputs

Our observations so far indicate moderate to poor accuracy of the networks inferred by the 12 methods on datasets from curated models. We were curious if the algorithms made similar errors or if there were some subnetworks of the Boolean models that were easy for most algorithms to infer correctly. We performed three analyses for each model separately. (i) We converted each algorithm’s output into a vector of edges and recorded their ranks in this vector. We used PCA on the set of vectors on all 10 datasets and all 12 algorithms to compute a two-dimensional projection (Supplementary Figure S9). (ii) For each dataset we computed the Spearman’s correlation coefficients between these vectors for all pairs of algorithms. Supplementary Figure S10 presents the results for one representative dataset. (iii) For the top-*k* edges for each dataset, we computed the Jaccard indices between all pairs of algorithms. Supplementary Figure S11 presents the results for one representative dataset. The PCA results suggest that some algorithms have somewhat similar outputs (e.g., GENIE3, GRNBoost2, PIDC, LEAP, and PPCOR in three out of the four models). The Spearman’s correlations and Jaccard indices echo these trends. GENIE3 and GRNBoost2 are both based on random forests, which may explain their similarities. Similarly, both LEAP and PPCOR are based on computing correlations. What is striking is that most pairs of Spearman’s correlations are somewhat negative (between −0.25 and 0) and most pairs of Jaccard indices are less than 0.5, suggesting that the outputs are quite different from each other. Notably, SCODE does not have much similarity to any other method.

## Discussion

We have presented BEELINE, a framework for benchmarking algorithms that infer GRNs from single-cell gene expression data. Although a few reviews of GRN inference methods have appeared [30–32], there has been only one quantitative evaluation of these approaches [21]. This paper compared eight approaches of which six were originally developed or use methods created for bulk transcriptomic data. In contrast, we include eight new algorithms that have been developed exclusively for single-cell gene expression data while not including some of the previously-compared approaches for reasons given in “Methods”. Most of the techniques we have considered have been developed since 2017. The earlier comparison used GeneNetWeaver to create simulated datasets. Thus, our evaluation is up-to-date and timely while also being more relevant for single-cell data.

We found considerable variation in the performance of the algorithms across the ten different networks (six synthetic and four Boolean) that we analyzed. Nevertheless, we noted a few trends. In general, the synthetic networks were easier to recover than the Boolean models. This trend may be due to the fact that the synthetic networks have simpler and more well-defined trajectories, which our simulations capture accurately, than the Boolean models. For datasets from curated models, each of which has multiple trajectories, we found that methods that do not require pseudotime information (specifically, GENIE3, GRNBoost2, and PIDC) performed the best. For the other techniques, we input pseudotime values for each trajectory independently. We found that these approaches succeeded in discovering mutual inhibition loops, with each arm of the loop inferred from a different trajectory. Nevertheless, the overall performance of these approaches was less than ideal. Note that three methods (GRISLI, SCINGE, and SCRIBE) have appeared only as preprints. Their final published versions may show improved performance.

Inspired by the success of ensembles in inferring GRNs from bulk transcriptional data [33], we attempted to combine the results of the algorithms we analyzed. The Borda method re-ranks every edge by the averages of its ranks from each GRN algorithm. We tried this approach and its variations, such as combining only the two or three methods with the highest AUROC, weighting all the methods by their AUROC [33], and down-weighting worse ranks [15]. To our surprise, none of these ensembles performed discernibly better than the algorithm with the highest AUROC for each synthetic network or Boolean model. This result suggests that the GRN algorithms are very diverse and perhaps even conflicting in their predictions, a conclusion borne out by our analysis of their similiarities (Supplementary Figures S9 to S11).

Despite the widely varying performance of GRN inference algorithms from one ground truth network to another, we can make some specific recommendations for users.

i. GRNBoost2 was a leading and consistent performer in our evaluations; it was among the four methods with the highest median AUROC value both for datasets from synthetic networks and for datasets from curated models. The results of this algorithm were also quite stable to variations in the input, at least for datasets from synthetic networks. It is conceivable that the random forest-based predictors that lie at its core may be further improved or specialized to single-cell contexts.
ii. LEAP, GENIE3, and GRNBoost2 may be suitable in scenarios where we do not expect too many feedforward loops in the true regulatory network.
iii. If efficiency is important, GRNBoost2, SINCERITIES, PPCOR and PIDC are preferable, since each of them had a running time of under a minute.
iv. More broadly, to determine which inference algorithm may be most appropriate for a specific experimental dataset, we suggest that a researcher apply our BoolODE-based approach: carefully select a Boolean or a dynamic (ODE-based) model of a relevant biological process [34], extract the underlying GRN, stochastically simulate this model to mimic a single-cell gene expression data set, invoke multiple GRN inference algorithms, and measure the accuracy of the outputs against the true GRN.

The final test of these algorithms is on how well they can predict GRNs when applied to experimentally-derived single-cell gene expression datasets. In the GRN inference literature, a common practice is to evaluate the accuracy of resulting network by comparing its edges to an appropriate database of TFs and their targets. For example, SCODE uses the RikenTFdb and animalTFDB resources to define ground truth networks for mouse and human gene expression data, respectively [6]. When we examined the results in the literature [9, 15], we found that the AUROC or precision at early recall values of these reconstructions were quite poor, often being close to that of a random predictor. Therefore, we reasoned that it would be more appropriate and useful to the community to benchmark GRN algorithms by applying them to carefully simulated Boolean models with predictable or computable trajectories.

In summary, our evaluation shows that GRN inference remains a challenging problem despite the variety of innovative ideas that researchers have brought to bear upon the problem. In this context, we obtained a striking finding that false positive edges form feedforward loops when added to ground truth networks. There is a history of GRN inference algorithms that explicitly try to avoid indirect effects [12, 35–37]. We suggest that new ideas for avoiding the prediction of such interactions may assist in improving the accuracy of GRN inference algorithms for single cell gene expression data. As single-cell experiments become more complex, we expect that transcriptional trajectories will also be more intricate, perhaps involving multiple stages of bifurcation and/or cycling. A key challenge that lies ahead is accurately computing GRNs that give rise to such complex trajectories. We hope that scientists in this field will use BEELINE in conjunction with BoolODE as they develop new approaches for GRN inference.

## Methods

### Regulatory Network Inference Algorithms

We briefly describe each algorithm we have included in this evaluation. We have ordered the methods chronologically by year and month of publication. Every software package had an open source licence, other than GRNVBEM, SCRIBE, and GRISLI, which did not have any licence.

1. **GENIE3 (GEne Network Inference with Ensemble of trees) [3].** Developed originally for bulk transcriptional data, GENIE3 computers the regulatory network for each gene independently. It uses tree-based ensemble methods such as Random Forests to predict the expression profile of each target gene from profiles of all the other genes. The weight of an interaction comes from the importance of an input gene in the predictor for a target gene’s expression pattern. Aggregating these weighted interactions over all the genes yields the regulatory network. This method was the top performer in the DREAM4 In Silico Multifactorial challenge.
2. **PPCOR [4].** This R package computes the partial and semi-partial correlation coefficients for every pair of variables (genes, in our case) with respect to all the other variables. It also computes a *p*-value for each correlation. We use this approach for the partial coefficients. Since these values are symmetric, this method yields an undirected regulatory network. We use the sign of the correlation, which is bounded between −1 and 1, to signify whether an interaction is inhibitory (negative) or activating (positive).
3. **LEAP (Lag-based Expression Association for Pseudotime-series) [5].** Starting with pseudotime-ordered data, LEAP calculates the Pearson’s correlation of normalized mapped-read counts over temporal window of a fixed size with different lags. The score recorded for a pair of genes is the maximum Pearson’s correlation over all the values of lag that the method considers. The software includes a permutation test to estimate false discovery rates. Since the correlation computed is not symmetric, this method can output directed networks.
4. **SCODE [6].** This method uses linear Ordinary Differential Equations (ODEs) to represent how a regulatory network results in observed gene expression dynamics. SCODE relies on a specific relational expression that can be estimated efficiently using linear regression. In combination with dimension reduction, this approach leads to a considerable reduction in the time complexity of the algorithm.
5. **PIDC (Partial Information Decomposition and Context) [7].** This method uses concepts from information theory. For every pair of genes *x* and *y*, given a third gene *z*, the method partitions the pairwise mutual information between *x* and *y* into a redundant and a unique component. It computes the ratio between the unique component and the mutual information. The sum of this ratio over all other genes *z* is the proportional unique contribution between *x* and *y*. The method then uses per-gene thresholds to identify the most important interactions for each gene. The resulting network is undirected since the proportional unique contribution is symmetric.
6. **SINCERITIES (SINgle CEll Regularized Inference using TIme-stamped Expression profileS) [9].** Given time-stamped transcriptional data, this method computes temporal changes in each gene’s expression through the distance of the marginal distributions between two consecutive time points using the Kolmogorov–Smirnov (KS) statistic. To infer regulatory connections between TFs and target genes, the approach uses Granger causality, i.e., it uses the changes in the gene expression of TFs in one time window to predict how the expression distributions of target genes shift in the next time window. The authors formulate inference as a ridge regression problem. They infer the signs of the edges using partial correlation analyses.
7. **SCNS [11].** This method takes single-cell gene expression data taken over a time course as input and computes logical rules (Boolean formulae) that drive the progression and transformation from initial cell states to later cell states. By design, the resulting logical model facilitates the prediction of the effect of gene perturbations (e.g., knockout or overexpression) on specific lineages.
8. **GRNVBEM [10].** This approach infers a Bayesian network representing the gene regulatory interactions. It uses a first-order autoregressive model to estimate the fold change of a gene at a specific time as a linear combination of the expression of the gene’s parents in the Bayesian network at the previous time point. It infers the gene regulatory network within a variational Bayesian framework. This method can associate signs with its directed edges but produces an acyclic graph (except for self-loops).
9. **SCRIBE [12].** This paper also uses ideas from information theory. The relevant concept here is conditioned Restricted Directed Information (cRDI), which measures the mutual information between the past state (expression values) of a regulator and the current state of a target gene conditioned on the state of the target at the previous time point. The authors subsequently use the Context Likelihood of Relatedness (CLR) algorithm [37] to remove edges that correspond to indirect effects.
10. **GRNBoost2 [13].** GRNBoost2 is a fast alternative for GENIE3, especially suited for datasets with tens of thousands of observations. Like GENIE3, GRNBoost2 trains a regression model to select the most important regulators for each gene in the dataset. GRNBoost2 achieves its efficiency by using stochastic Gradient Boosting Machine regression with early-stopping regularization to infer the network.
11. **GRISLI [14].** Like SCODE, this approach uses a linear ODE-based formalism. GRISLI estimates the parameters of the model using different ideas. Taking either the experimental time of the cells or estimated pseudotime as input, it first estimates the velocity of each cell, i.e., how each gene’s expression value changes as each cell undergoes a dynamical process [38]. It then computes the structure of the underlying gene regulatory network by solving a sparse regression problem that relates the gene expression and velocity profiles of each cell.
12. **SCINGE (Single-Cell Inference of Networks using Granger Ensembles) [15].** The authors observe that while many gene inference algorithms start by computing a pseudotime value for each cell, the distribution of cells along the underlying dynamical process may not be uniform. To address this limitation, SCINGE uses kernel-based Granger Causality regression to alleviate irregularities in pseudotime values. SCINGE performs multiple regressions, one for each set of input parameters, and aggregates the resulting predictions using a modified Borda method.

#### Other methods

We now discuss other papers on this topic and our rationale for not including them in the comparison. We did not consider a method if it was supervised [39] or used additional information, e.g., a database of transcription factors and their targets [8], a lineage tree [17]. We did not include methods that output a single GRN without any edge weights [18, 23], since any such approach would yield just a single point on a precision-recall or curve.

## Datasets

### Creating Datasets from Synthetic Networks

We now describe how we selected synthetic networks, converted these networks into systems of stochastic differential equations (SDEs), simulated these systems, and pre-processed the resulting datasets for input to GRN inference algorithms.

#### Selecting networks

In order to create synthetic datasets exhibiting diverse temporal trajectories, we use six “toy” networks created by Saelens *et al.* [22] in their comparison of pseudotime inference algorithms (Figure 2 and Table 1). When simulated as SDEs, we expect the models produce to trajectories with the following qualitative properties:

**Linear:** a gene activation cascade that results in a single temporal trajectory with distinct final and initial states.
**Linear long:** similar to Linear but with a larger number of intermediate genes.
**Cycle:** an oscillatory circuit that produces a linear trajectory where the final state overlaps with the initial state.
**Bifurcating:** a network that contains a mutual inhibition motif between two genes resulting in two distinct branches starting from a common trajectory.
**Trifurcating:** three consecutive mutual inhibition motifs in this network result in three distinct steady states.
**Bifurcating Converging:** an initial bifurcation creates two branches, which ultimately converge to a single steady state.

**Table 1:** Summary of synthetic networks. An asterisk indicates limit cycle oscillations.

| Name of network | Number of genes | Number of edges |  | Number of Steady States |
| --- | --- | --- | --- | --- |
|  |  | Activation | Inhibition |  |
| Linear | 7 | 7 | 1 | 1 |
| Linear Long | 18 | 18 | 1 | 1 |
| Cycle | 6 | 3 | 3 | 1* |
| Bifurcating | 7 | 7 | 5 | 2 |
| Trifurcating | 8 | 10 | 10 | 3 |
| Bifurcating Converging | 10 | 13 | 5 | 1 |

We do not consider three other networks (Consecutive Bifurcating, Bifurcating Loop and Converging) created by Saelens *et al.* since we were unable to recapture the expected trajectories using the approach described below.

#### Converting networks into SDE models and simulating them

We manually convert each of these networks to a Boolean model: we set a node to be ‘on’ if and only if at least one activator is ‘on’ and every inhibitor is ‘off’. We simulate these networks using BoolODE, a method we have developed to convert Boolean models into systems of differential equations (Supplementary Section S1). We use the initial conditions specified in the Dynverse software created by Saelens *et al.* (Supplementary Table S2). In order to sample cells from the simulations that capture various locations along a trajectory, we limit the duration of each simulation according to the characteristics of the model (Table S2). For example, we simulated the Linear Long network for 15 steps but the Linear network, which has fewer nodes, for 5 steps.

#### Comparing simulated datasets with the expected trajectories from synthetic networks

It is common to visualize simulated time courses from ODE/SDE models as time course plots or phase plane diagrams with two or three dimensions. The latter are useful to qualitatively explore the state-space of a model at hand. The recent popularity of t-SNE as a tool to visualize and cluster high dimensional scRNAseq data motivated us to consider this technique to visualize simulated ‘single cell’ data. Figure 2 shows two-dimensional t-SNE visualizations of each of the toy models. The color of each point on the t-SNE scatter plot reflects the corresponding simulation timepoint (blue for early, green for intermediate, and yellow for later time points). We find that the t-SNE visualizations succeed in capturing the qualitative steady states of each of the models we simulate. Specifically, the number of steady state clusters we expect from each synthetic network (e.g., one steady state for linear, two for bifurcating, etc.) are captured with great fidelity in the t-SNE visualizations.

#### Pre-processing datasets from synthetic networks for GRN algorithms

Saelens *et al.* [22] have shown that pseudotime inference methods perform well for linear and bifurcating trajectories. However, even the best performing pseudotime algorithm fails to accurately identify more complex trajectories such as Cycle and Bifurcating Converging. Therefore, we sought to develop an approach for pre-processing synthetic datasets that mimicked a real single cell gene expression pipeline while isolating the GRN inference algorithms from the limitations of pseudotime techniques. Accordingly, we used the following four-step approach to generate single-cell gene expression data from each of the six synthetic networks:

1. Using the parameters in Table S2, we performed 150 simulations, sampling 100 cells per simulation, thus obtaining a total of 15,000 cells.
2. We represent each simulation as a |*G*| × |*C*| matrix, where *G* is the set of genes in the model and *C* is the set of cells in the simulation. We convert each of the 150 matrices into a 1-D vector of length |*G*| × |*C*| and cluster the vectors using *k*-means clustering with *k* set to the number of expected trajectories for each network. For example, the Bifurcating network in Figure 2 has two distinct trajectories, so we used *k* = 2.
3. We then randomly sampled 10 different sets of 500, 2000, and 5000 cells from the 15,000 cells in such a way that there were an equal number of cells sampled from each individual trajectory.
4. Finally, we set as input to each algorithm the |*G*| × |*C*| matrix, where *G* is the set of genes in the model and C is the set of randomly sampled cells. For those methods that require time information, we specified the simulation time at which the cell was sampled. Thus, we ran each algorithm on the same set of 30 datasets.

### Creating Datasets from Curated Models

While the synthetic models presented above are useful for generating simulated data with a variety of specific trajectories, these networks do not correspond to any real cellular process. In order to create simulated datasets that better reflect the characteristics of single-cell transcriptomic datasets, we turned to published Boolean models of gene regulatory networks, as these models are reflective of the real “ground truth” control systems in biology.

Since tissue differentiation and development are active areas of investigation by single cell methods, we examined the literature from the past 10 years to look for published Boolean models of gene regulatory networks involved in these processes. We selected four published models for analysis. Table 2 lists the size of the regulatory networks and the number of steady states. Below, we discuss the biological background, the interpretation of model steady states, and the expected type of trajectories for each of these Boolean models.

**Table 2:** Summary of published Boolean models.

| Name of model | Number of genes | Number of edges |  | Number of Steady States | Citation |
| --- | --- | --- | --- | --- | --- |
|  |  | Activation | Inhibition |  |  |
| mCAD | 5 | 5 | 9 | 2 | [25] |
| VSC | 8 | 0 | 15 | 5 | [26] |
| HSC | 11 | 15 | 15 | 4 | [27] |
| GSD | 19 | 27 | 59 | 2 | [28] |

#### Mammalian Cortical Area Development

Giancomantonio *et al.* explored mammalian Cortical Area Development (mCAD) as a consequence of the expression of regulatory transcription factors along an anterior-posterior gradient [25]. The model contains five transcription factors connected by 14 interactions, captures the expected gene expression patterns in the anterior and posterior compartments respectively, and results in two steady states. Figure 4 displays the regulatory network underlying the model along with a t-SNE visualization of the trajectories simulated using BoolODE. Figure S3 shows that the two clusters observed in the t-SNE visualization correspond to the two biological states captured by the Boolean model.

#### Ventral Spinal Cord Development

Lovrics *et al.* investigated the regulatory basis of ventral spinal cord development [26]. The model consisting of 8 transcription factors involved in ventralization contains 15 interactions, all of which are inhibitory. It succeeds in accounting for five distinct neural progenitor cell types. We expect to see five steady states from this model. Figure 4 shows the regulatory network underlying the model along with the t-SNE visualization of the trajectories simulated using BoolODE. Supplementary Figure S4 shows that the five steady state clusters observed in the t-SNE visualization correspond to the five biological states captured by the model.

#### Hematopoietic Stem Cell Differentiation

Krumsiek *et al.* investigated the gene regulatory network underlying myeloid differentiation [27]. The proposed model has 11 transcription factors and captures the differentiation of multipotent Myeloid Progenitor (CMP cells) into erythrocytes, megakaryocytes, monocytes and granulocytes. The Boolean model exhibits four steady states, each corresponding to one of the four cell types mentioned above. Figure 4 shows the regulatory network of the HSC model along with the t-SNE visualization of the trajectories simulated using BoolODE. Supplementary Figure S5 shows that the four steady-state clusters observed in the t-SNE visualization correspond to the four biological states captured by the Boolean model.

#### Gonadal Sex Determination

Rios *et al.* 2015 modeled the gonadal differentiation circuit that regulates the maturation of the Bipotential Gonadal Primordium (BGP) into either male (testes) or female (ovary) gonads [28]. The model consists of 18 genes and a node representing the Urogenital Ridge (UGR), which serves as the input to the model. For the wildtype simulations, the Boolean model predominantly exhibits two steady states corresponding to the Sertoli cells (male gonad precursors) or the Granulosa cells (female gonad precursor), and one rare state corresponding to a dysfunctional pathway. Figure 4 shows the regulatory network of the GSD model along with the t-SNE visualization of the trajectories simulated using BoolODE. Supplementary Figure S6 shows that the two steady state clusters observed in the t-SNE visualization correspond to the two predominant biological states capture by the Boolean model.

#### Pre-processing datasets from curated models for GRN algorithms

We simulated the four curated models using parameters mentioned in Supplementary Table S2. As with the datasets from synthetic networks, we performed 150 simulations, and sampled 100 cells per simulation for a total of 15,000 cells. For each model, we then randomly sampled ten different datasets, each containing 2,000 cells. In order to closely mimic the preprocessing steps performed for real single-cell gene expression data, we use the following steps for each dataset:

1. *Pseudotime inference using Slingshot*. In the case of datasets from synthetic networks, we used the simulation time directly for those GRN inference methods that required cells to be time-ordered. In contrast, for datasets from curated models data, we ordered cells by pseudotime, which we computed using Slingshot [29]. We selected Slingshot for pseduotime inference due to its proven success in correctly identifying cellular trajectories in a recent comprehensive evaluation of this type of algorithm [22]. Slingshot needs a lower dimensional representation of the gene expression data as input. In addition, if the cells belong to multiple trajectories, Slingshot needs a vector of cluster labels for the cells, as well as the cluster labels for cells in the start and end states in the trajectories. To obtain these additional input data for Slingshot, we used the following procedure:

a. We use t-SNE on the |*G*| × |*C*| matrix representing the data to obtain a two-dimensional representation of the cells.
b. We performed *k*-means clustering on the lower dimensional representation of the cells with *k* set to one more than the expected number of trajectories. For example, since we knew that the GSD model had a bifurcating trajectory, we used *k* = 3.
c. We computed the average simulation time of the cells belonging to each of the k clusters. We set the cluster label corresponding to the smallest average time as the starting state and the rest of clusters as ending states.
d. We then ran Slingshot with the t-SNE-projected data and starting and ending clusters as input. We obtained the trajectories to which each cell belonged and its pseudotime as output from Slingshot. Figure 4 display the results for one dataset of 2,000 cells for each of the four model. We observed that Slingshot does a very good job of correctly identifying cells belonging to various trajectories. Further, the pseudotime computed for each cell by Slingshot is highly correlated with the simulation time at which the cell was sampled (data not shown).
2. *Inducing dropouts in datasets from curated models*. We used the same procedure as Chan *et al.* [7] in order to induce dropouts, which are commonly seen in single cell RNA-Seq datasets, especially for transcripts with low abundance [40]. We applied two different dropout rates: *q* = 20 and *q* = 50. For every gene, we sorted the cells in increasing order of that gene’s expression value. We set the expression of that gene in each of the lowest *q^th^* percentile of cells in this order to zero with a 50% chance. We did not recompute pseudotime for these new datasets with dropouts. Instead, we used the pseudotime values computed on the dataset without dropouts.

At the end of this step, we had 30 datasets with 2,000 cells for each of the four models: 10 datasets without dropouts, 10 with dropouts at *q* = 20, and 10 with dropouts at *q* = 50.

### Evaluation Pipeline

One of the major challenges we faced was that the GRN inference methods we included in this evaluation were implemented in a variety of languages such as R, MATLAB, Python, Julia and F#. In order to obtain an efficient and reproducible pipeline, we Dockerized the implementation of each of these algorithm. Supplementary File 1 contains details on the specific software or GitHub commit versions we downloaded and used in our pipeline. For methods implemented in MATLAB, we created MATLAB executable files (.mex files) that we could execute within a Docker container using the MATLAB Runtime. We further studied the publications and the documentation of the software (and the source code, on occasion) to determine how the authors recommended that their methods be used. We have incorporated these suggestions as well as we could. We provide more details in Supplementary Section S3.

#### Inputs

For datasets from synthetic networks and for datasets from curated models, we provided the gene expression values obtained from the simulations directly after optionally inducing dropouts in the second type of data. As shown in Figure 3, eight out of the 12 methods also required some form of time information for every cell in the dataset. Of these, two methods (GRNVBEM and LEAP) only required cells to be ordered according to their pseudotime and did not require the pseudotime values themselves.

#### Parameter estimation

Six of the methods also required one or more parameters to be specified. To this end, we performed parameter estimation for each of these methods separately for datasets from synthetic networks, datasets from curated models, and real datasets and provided them with the parameters that resulted in the best AUROC values. See Supplementary Section S2 for details.

#### Output processing

Ten of the GRN inference methods output a confidence score for every possible edge in the network, either as an edge list or as an adjacency matrix, which we converted to a ranked edge list. We gave edges with the same confidence score the same rank.

#### Performance evaluation

We used a common evaluation pipeline across all the datasets considered in this paper. We evaluated the result of each algorithm using the following criteria:

1. **AUROC, AUPRC**: We computed areas under the Receiver Operating Characteristic (ROC) and Precision-Recall (PR) curves using the edges in the relevant ground truth network as ground truth and ranked edges from each method as the predictions. We ignored self-loops for this analysis since some methods such as PPCOR always provide the highest rank for such edges and some other methods such as SCINGE and always ignore them.
2. **Identifying top-*k* edges**: We first identified top-*k* edges for each method, where *k* equaled the number of edges in the ground truth network (excluding self loops). In multiple edges were tied for a rank of *k*, we considered all of them. If a method provided a confidence score for fewer than *k* edges, we used only those edges.
3. **Jaccard index**: Once we obtained the set of top-*k* edges for each method and each dataset, we then computed the Jaccard index of every pair of these sets. We used the median of the values as an indication of the robustness of a method’s output to variations in the simulated datasets from a given synthetic network or Boolean model.
4. **Early precision**: We defined early precision as the fraction of true positives in the top-*k* edges. We also computed early precision ratio, which represents the ratio of early precision value and the early precision for a random predictor for that model. A random predictor’s precision is the edge density of the ground truth network.
5. **Early precision of signed edges**: We desired to check whether there were any differences in how accurately a GRN inference algorithm identified activating edges in comparison to inhibitory edges. To this end, we computed the top-*k_a_* edges from the ranked list of edges output by each method, where *k_a_* is the number of activating edges in the ground-truth network. In this step, we ignored any inhibitory edges in the ground truth network. We defined the early precision of activating edges as the fraction of true edges of this type in the top *k_a_* edges. We used an analogous approach to compute early precision of inhibitory edges. We also computed the early precision ratio for these values.
6. **Network motifs**: In each proposed network formed using the top-*k* predicted edges, we counted the number of three-node feedback loops, feedforward loops and mutual interactions involving every edge and divided this number by their corresponding values in the ground truth networks. For this analysis, we ignored algorithms that predict only undirected edges (PIDC and PPCOR).
7. **Connectivity of false positive edges**: We were interested in determining if GRN methods were successful in weeding out indirect interactions from their predictions. To this end, for each false positive edge in a top-*k* network, we compute the length of the shortest path between the endpoints of that edge in the relevant ground truth network. We plotted histograms of the distributions of these distances. We ignored algorithms that predict only undirected edges for this analysis as well.

#### Datasets with multiple trajectories

Most methods cannot directly handle data with branched trajectories. In such situations, as suggested uniformly by many of these methods, we separate the cells into multiple linear trajectories using Slingshot, the top performing pseudotime algorithm in a recent comparison [22], and apply the algorithm to the set of cells in each trajectory individually. To combine the GRNs, for each interaction, we record the largest score for it across all the networks, and rank the interactions by these values. In the case of GRISLI, which outputs ranked edges, for each interaction, we took the best (smallest) rank for it across all the networks.

## Declarations

### Availability of data and material

A Python implementation of our complete framework is available under the GNU General Public License v3 at https://github.com/murali-group/BEELINE

## Acknowledgements

Grants from the National Science Foundation (CCF-1617678, DBI-1759858) and the National Cancer Institute (UH2CA203768) supported this work. The research is also based upon work supported by the Office of the Director of National Intelligence (ODNI), Intelligence Advanced Research Projects Activity (IARPA), via the Army Research Office (ARO) under cooperative Agreement Number [W911NF-17-2-0105]. The views and conclusions contained herein are those of the authors and should not be interpreted as necessarily representing the official policies or endorsements, either expressed or implied, of the ODNI, IARPA, ARO, or the U.S. Government. The U.S. Government is authorized to reproduce and distribute reprints for Governmental purposes notwithstanding any copyright annotation thereon. The funding bodies played no role in the design of the study, the collection, analysis, and interpretation of data, or in writing the manuscript.

## Authors’ contributions

AP studied the literature, selected the GRN algorithms, implemented the pipeline, led the analysis, and wrote the paper. APJ implemented BoolODE, created the datasets from synthetic networks and datasets from curated models, selected the real gene expression datasets, and wrote the paper. JNL performed and described the parameter searches. AB created network layouts, performed the Borda and comparative analysis. TMM supervised the project, provided input on the GRN algorithms, BoolODE, and evaluation strategies, and wrote the paper.

## Supplementary Text

### S1 BoolODE: Converting Boolean Models to Ordinary Differential Equations

We start this section by giving an overview of GeneNetWeaver [24, 41], a popular method for simulating bulk gene expression datasets. It is being used increasingly for simulating single cell transcriptional data as well [7, 9, 10, 18, 21]. Next, we describe the BoolODE framework that we have developed and highlight its differences with GeneNetWeaver. We compare the data simulated by BoolODE and GeneNetweaver to demonstrate that the latter approach does not yield expected trajectories. We end this section by summarising BoolODE and the reasons we prefer it over GeneNetWeaver.

#### GeneNetWeaver

This method starts with a network of regulatory interactions among transcription factors (TFs) and their targets. It computes a connected, dense subnetwork around a randomly selected seed node and converts this network into a system of differential equations. To express this network in the form of Ordinary Differential Equations (ODEs), it assigns each node *i* in the network a ‘gene’ variable (*x_i_*) representing the level of mRNA expression and a ‘protein’ variable (*p_i_*) representing the amount of transcription factor produced by protein translation as follows:

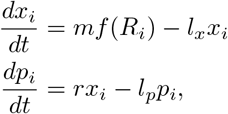

where *m* is the mRNA transcription rate, *l_x_* is the mRNA degradation rate, *r* is the protein translation rate, and *l_p_* is the protein degradation rate. In the first equation, *R_i_* denotes the set of regulators of node *i*. The non-linear input function *f*(*R_i_*) captures all the regulatory interactions controlling the expression of node *i* [20].

If there are *N* regulators for a given gene, there are 2*^N^* possible configurations of how the regulators can bind to the gene’s promoter. Considering cooperative effects of regulator binding, the probability of each configuration *S* ∈ 2^*R_i_*^, the powerset of *R_i_*, is given by the following equation [42]:

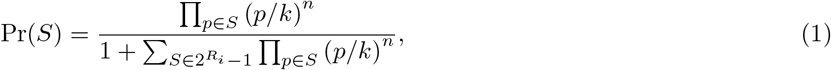

where *k* and *n* are the Hill threshold and Hill coefficient respectively. Here the product in the numerator ranges over all regulators that are present (bound) in a specific configuration *S*, and the sum in the denominator runs over all configurations in the powerset, other than the empty set. GeneNetWeaver further introduces a randomly-sampled parameter *α_S_* ∈ [0, 1] to specify the efficiency of transcription activation by a specific configuration S of bound regulators. Thus, the function *f* (*R_i_*) thus takes the following form:

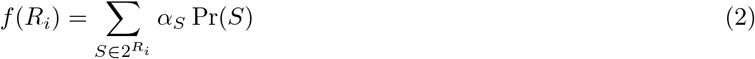

Next, GeneNetWeaver adds a noise term to each equation to mimic stochastic effects in gene expression [24]. In addition, in order to create variations among individual experimental samples, GeneNetWeaver recommends adopting a multifactorial perturbation [24] that increases or decreases the basal activation of each gene in the GRN simultaneously by a small, randomly-selected value. GeneNetWeaver removes this perturbation after the first half of the simulation. Simulating this system of SDEs generates the requisite gene expression data.

#### BoolODE uses Boolean models to create simulated datasets

In order to generate simulated time course data for our analysis, we used the GeneNetWeaver framework with one critical difference and one minor variation. The form of the equations used by BoolODE is identical to that of GeneNetWeaver. The critical difference is that we do not sample the *α_S_* parameters in Equation 2 randomly, i.e., we do not combine the regulators of each gene using a random logic function. Instead, we use the fact that in both the artificial networks and the literature-curated models, we know the Boolean function that specifies how the states of the regulators control the state of the target genes. Moreover, we can express any arbitrary Boolean function in the form of a truth table relating the input states (i.e., activities of transcription factors) to the output state (the activity of target gene). For a gene with *N* regulators in its Boolean function, we explore all 2*^N^* combinations of transcription factor states and evaluate the transcriptional activity of each specific regulator configuration. Since the value of the Boolean function is the logical disjunction (‘or’) of all these values, we set the *α* value to one (respectively, zero) for every configuration that evaluates to ‘on’(respectively, ‘off’). The following example illustrates our approach. Consider a gene *X* with two activators (*A* and *B*) and one inhibitor (*C*), represented by the following rule:

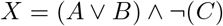

The truth table corresponding to this rule along with the *α* parameters appears below.

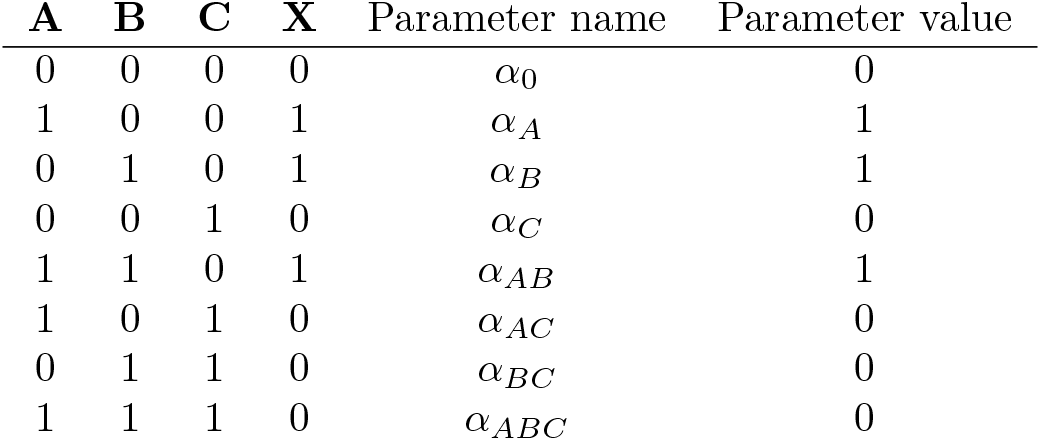

Therefore, the ODE governing the time dynamics of gene *X* is

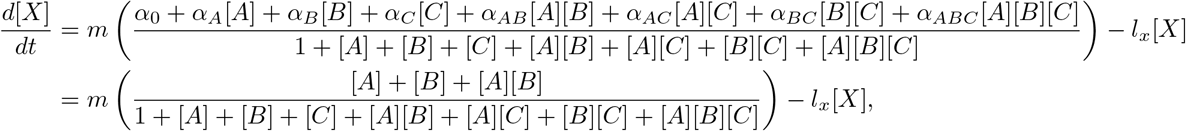

since only *α_A_*, *α_B_*, and *α_AB_* have the value one and every other parameter has the value zero.

Next, we discuss the minor variation of BoolODE from GeneNetWeaver, which is in how we sample kinetic parameters. The GeneNetWeaver equations use four kinetic parameters: one each for mRNA transcription, protein translation, and mRNA and protein degradation rates. Saelens *et al.* [22] sample them uniformly from parameter-specific intervals. We reasoned that a random sampling of parameters is undesirable as we wish to get reproducible temporal trajectories for each model that we consider. Therefore, we fix the kinetic parameters in our simulations. We set them to be the upper limits of the ranges used by Saelens *et al.* [22] (see Table S1 for the values we use).

**Table S1:** Kinetic parameters used in BoolODE.

| Parameter | Symbol | value |
| --- | --- | --- |
| mRNA transcription rate | $m$ | 20 |
| mRNA degradation rate | $l_x$ | 10 |
| Protein translation rate | $r$ | 10 |
| Protein degradation rate | $l_p$ | 1 |
| Hill threshold | $k$ | 10 |
| Hill coefficient | $n$ | 10 |

In order to create stochastic simulations, we use the formulation proposed by Saelens *et al.* [22]. We modify the ODE expressions as follows:

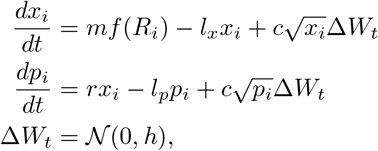

where *c* is the noise strength. We use *c* = 10 in our simulations. We use the Euler-Maruyama scheme for numerical integration of the SDEs with time step of *h* = 0.01.

#### Defining a single cell

We define the vector of gene expression values corresponding to a particular time point in a model simulation as a single cell. For all analysis, we sample 100 cells at uniform timepoints from every simulation.

#### Creating GeneNetWeaver simulations for comparison with BoolODE

In order to simulate a synthetic network using GeneNetWeaver, we used its the edge list as the input network to GeneNetWeaver. In order to create the simulations, we used the default options of noise parameter (0.05) and multifactorial perturbations. We only performed wildtype simulations and used the DREAM4 time series output format for comparison with the BoolODE output.

#### Summary

We developed the BoolODE approach to convert the Boolean functions directly to ODE equations. Our proposed BoolODE pipeline accepts a file describing a Boolean model as input, creates an equivalent ODE model, adds noise terms, and numerically simulates a stochastic time course. Different model topologies can produce different numbers of steady states. Since we carry out stochastic simulations, we perform a large number of simulations in an attempt to ensure that we can reach every steady state. Our earlier analysis of the trajectories computed by BoolODE on datasets from curated models demonstrates the success of our approach in this regard. We prefer BoolODE over a direct application of GeneNetWeaver to create datasets from synthetic networks and datasets from curated models for three reasons: (a) A dense regulatory subnetwork computed around a randomly-selected node, as used by GeneNetWeaver, may not correspond to a real biological process. (b) GeneNetWeaver introduces a random, initial, multifactorial perturbation and removes it halfway in order to create variations in the expression profiles of genes across samples. This stimulation may not correspond to how single-cell gene expression data is collected. (c) GeneNetWeaver’s SDEs do not appear to capture single-cell expression trajectories, as we have shown in Figure 2.

### S2 Parameter Search

Six of the 12 algorithms we tested had no parameters. For each of the other six methods, we performed a parameter sweep. We optimized the parameters on the datasets from the Linear synthetic network with three different numbers of cells (500, 2,000 and 5,000) and used the same parameters for the other synthetic networks. In the case of the four datasets from curated models, we ran each algorithm on each individual trajectory for each dataset, combined the ranked lists of edges as discussed in the main text, and computed the AUROC. We chose this approach since every GRN inference algorithm in our evaluation that had parameters and had been developed explicitly for single-cell gene expression data could take only one trajectory as input. Since each of these datasets had 10 replicates, we chose the parameter value or combination of values that maximized the median AUROC for each method (Supplementary Figures S12 and S13). Below we describe the parameters tested for each method. We also provide brief comments on the effects of the parameters on the AUROC.

#### LEAP

This method has a single parameter, `maxLag`, which controls the maximum amount of lag to consider. The authors of this method suggest not using a value larger than 1/3. Hence, we tested seven values in the interval [0, 0.33]: 0.05, 0.1, 0.15, 0.2, 0.25, 0.3, and 0.33.

#### SCODE

While SCODE has two parameters,`D`, and the number of iterations, we only varied the parameter `D`. We used the same value of 100 as the authors. We also experimented with 1, 000 iterations but found that this value did not improve the AUROC (data not shown). In the documentation for the SCODE code, the authors suggest averaging the results from multiple runs. Therefore, we averaged the results across five runs. For `D`, we tested the values that they evaluated in their paper: 2, 4, 6, 8, and 10. We found this parameter had a large effect on the AUROC and thus for each dataset, we used the value that gave the highest AUROC.

#### SINCERITIES

SINCERITIES does not have any direct parameters. However, since it was developed to utilize temporal transcriptional data, rather than pseudotime, we opted to discretize the pseudotime values into bins to represent time points. We varied the number of bins from 5 to 20 with the specific values of 5, 6, 7, 8, 9, 10, 15, and 20.

#### SCRIBE

In their preprint, they used the combination “5,10,20,25” for a parameter called delay. We evaluated this value as well as several other combinations of delays: “5”, “5,10”, “5,10,15,20”, “5,10,15,20,25”, “5,10,20,25”, “5,10,25,50,75,100”, and “20,25,50,75,100”.

#### GRISLI

This method has three parameters: `R, L`, and α. The authors suggest that `R` be as large as possible to reduce random fluctuations of the algorithm. We chose to use *R* = 3,000. Following the authors’ recommendation to test `L` and α over a large grid, we tried values in the same ranges as in their paper: `L` = 1, 5, 10, 25, 50, 75, and 100 and α= 0, 0.1, 0.2, 0.3, 0.4, 0.5, 0.6, 0.7, 0.8, 0.9, and 1.0.

#### SCINGE

SCINGE has a total of seven parameters. We tested the same ranges of parameter values as the authors tested in their preprint: sparsity λ = 0, 0.01, 0.02, 0.05, 0.1; *δt* and *L* combinations of (3,5), (5,9), (9,5), (5,15), (15,5); kernel width *σ* = 0.5, 1, 2, 4; probability of zero removal and probability of removing samples values = 0, and 0.2 each; and the number of replicates = 2, 6, and 10. These parameter values resulted in a total of 1200 combinations. While the authors of SCINGE used an ensemble of parameter values, we chose to simply use the combination of parameters that resulted in the best AUROC. Since varying the parameters corresponding to the number of replicates, probability of zero removal and the probability of removing samples did not affect the AUROC, we chose 6, 0 and 0 for their values. For λ, small values of 0 or 0.01 were best. *δt*, *L* and *σ* had the largest effect on AUROC.

### S3 Software Infrastructure

In this section, we provide more information on how we executed each algorithm.

#### GENIE3 and GRNBoost2

We used the Python package Arboreto [13] for running GENIE3 and GRNBoost2. We choose this implementation of GENIE3 since this is the version used in an application of this method on single-cell data [8]. The current implementation of GENIE3 and GRNBoost2 in Arboreto takes an expression matrix and a list of genes as input. It produces a ranked list of edges as output, which we used directly in our analysis.

#### PPCOR

We used the R implementation of PPCOR available on CRAN. The software has options to use non-parametric partial and semi-partial correlation coefficients based on Kendall’s and Spearman’s rank correlations. We used Spearman’s correlation. We used a cut-off of 0.01 on the *p*-values computed by PPCOR to decide whether to include an interction in the output or not, similar to the approach used in Pseudotime Network Inference [19]. We ordered the interactions in decreasing order of absolue value of the correlation.

#### LEAP

We used the R implementation of LEAP. The software has options for cutoffs for maximum lag and maximum absolute correlation (MAC) and whether or not to obtain a directed network as output. We choose the option to obtain a directed network. Since we were interested in obtaining a rank for every edge, we set the MAC cutoff to 0. We then performed a parameter search on the maximum lag as described in the “Parameter Search”section. LEAP outputs an adjacency matrix, which we converted to a ranked list of edges based on the MAC score.

#### SCODE

SCODE is available both as an R and a Julia package. We used the R implementation. The software has options for the number of reduced dimensions, number of iterations for optimizing ODE parameters, and for running SCODE several times. We describe the parameter search for SCODE in the “Parameter Search” section. SCODE outputs an adjacency matrix *A*, with positive and negative values for edge weights indicating activation or inhibition. In order to obtain the ranked list of edges, we sorted the edges by the absolute values of the edge weights.

#### PIDC

PIDC is available in a Julia package titled “NetworkInference”. This package has options to compute various network inference algorithms such has Mutual Information, CLR, PUC and PIDC [7]. We used the maximum likelihood estimator, as recommended by the authors. We did not use any edge-weight cut-off. We converted the resulting undirected network in the form of an adjacency matrix into to a ranked edge list.

#### SINCERITIES

SINCERITIES is available both as an R and a MATLAB package. We used the R implementation. The software has options for the distance parameter, regularization strategy, and whether or not to obtain self loops. We used the Kolmogorov-Smirnov distance and Ridge regularization as described in the SINCERITIES paper. SINCERITIES also recommends binning pseduotime values into a pre-specified number of bins. Therefore, we performed a parameter estimation on number of bins. SINCERITIES outputs a ranked list of edges along with edge weights, which we used directly in our analysis.

#### SCNS

We used the F# implementation of SCNS. SCNS requires the following inputs: binarized gene expression values for each cell, a list of cells in the initial state, a list of cells in final states, maximum numbers of activators and inhibitors for each gene, and a threshold parameter. We first binarized the gene expression data by setting the gene’s expression to zero if it is lower than average expression of that gene across all cells. We sorted the cells by pseudotime and specified that the cells in the bottom 10th percentile were in the initial state and those in the top 10th percentile were in the final state. For each gene we specified the same number of activators and inhibitors as in the reference network, with a threshold parameter of 100. For each gene, this method outputs a Boolean formula that combines the states (on/off) of the regulators to produce the state of the regulated gene. The formula has a specific structure: the gene is ‘on’ if and only at least one activator is ‘on’ and all inhibitors are ‘off’. We convert each such formula into a directed network as follows: we create an edge directed from each regulator to the target gene. For our analysis we ignored whether a regulator acts as an activator or inhibitor, and simply created an edge if the regulator appears in the rule. For each gene, SCNS also outputs multiple Boolean formulae that satisfy the given input parameters. Therefore, if a regulator appeared in multiple rules, we counted the number of rules in which a regulator appears and used this quantity as the edge weight. We sorted this list of edges based on their weights to obtain the ranked edge lists used in our analysis.

#### SCRIBE

SCRIBE is available as an R package. Among the many variants of Restricted Directed Information presented in this paper, we used the ucRDI with *k* =1, since this alternative appeared to have the highest accuracy; *k* = 1 means computing the RDI value conditioned on each other gene individually in order remove indirect causal interactions The paper says that after applying CLR, it is possible to further sparsify the network using a method for regularization. However, we did not include this step since the documentation for their R package does not give any guidelines on what functions to invoke here. SCRIBE outputs an adjacency matrix of RDI values, which we then converted to an edge list.

#### GRISLI

GRISLI is available as a MATLAB package. GRISLI has options for parameters such as *R, L* and *α*, for which we performed parameter estimation using values recommended by the authors. GRISLI outputs a list of ranks for each edge as an adjacency matrix, which we then converted to a ranked edge list.

#### SCINGE

SCINGE is available both as a MATLAB package. The software has options for various parameters, for which we performed parameter estimation. SCINGE outputs a ranked list of edges along with edge weights, which we used directly in our analysis.

### S4 Supplementary Figures

**Figure S1:**
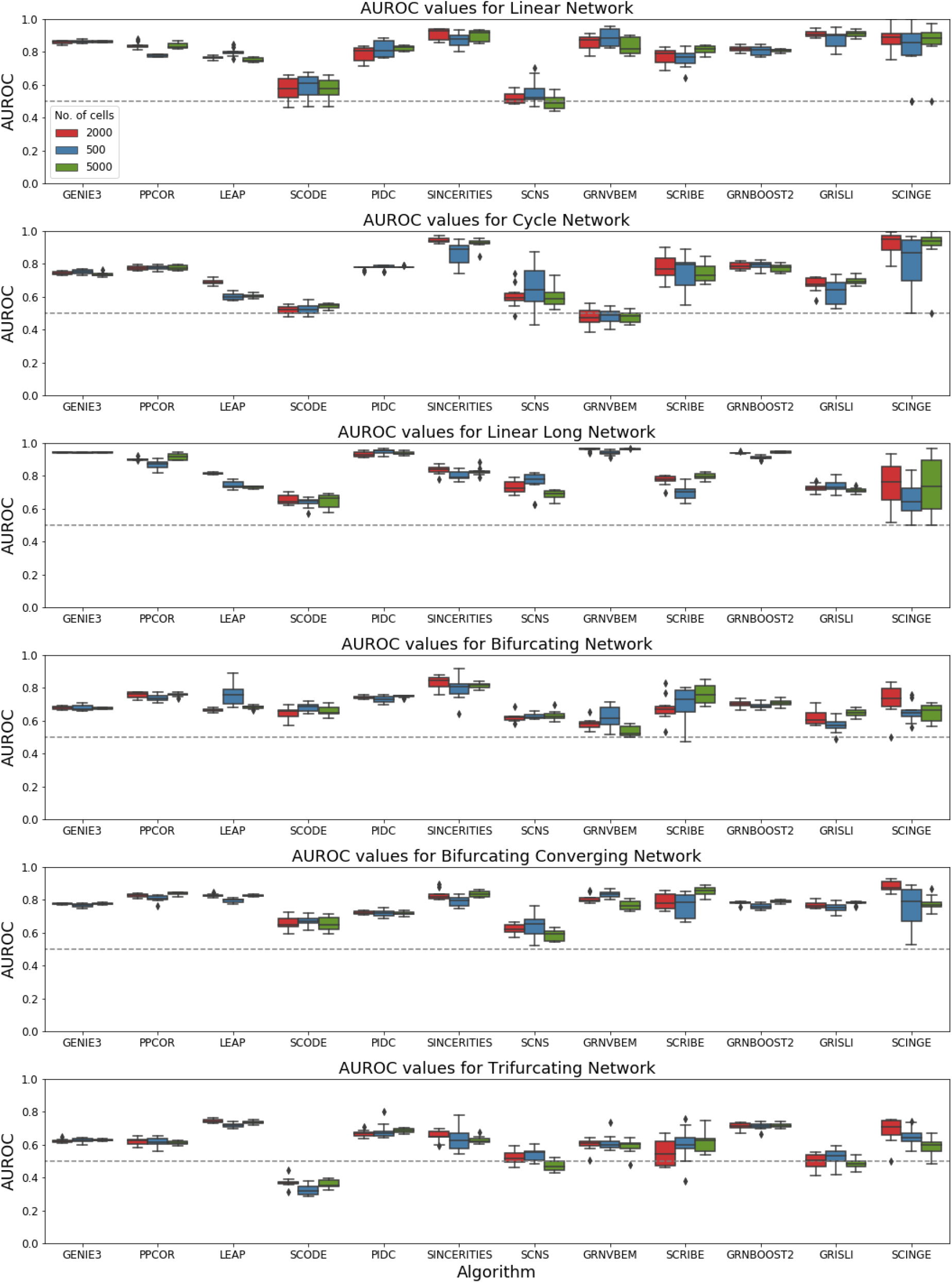
Box plots of AUROC values. Each row corresponds to one of the six synthetic networks. Each column corresponds to an algorithm. Blue, red, and green box plots corresponds to datasets with 500, 2,000, and 5,000 cells, respectively. The gray dotted line indicates the AUROC value for a random predictor (0.5).

**Figure S2:**
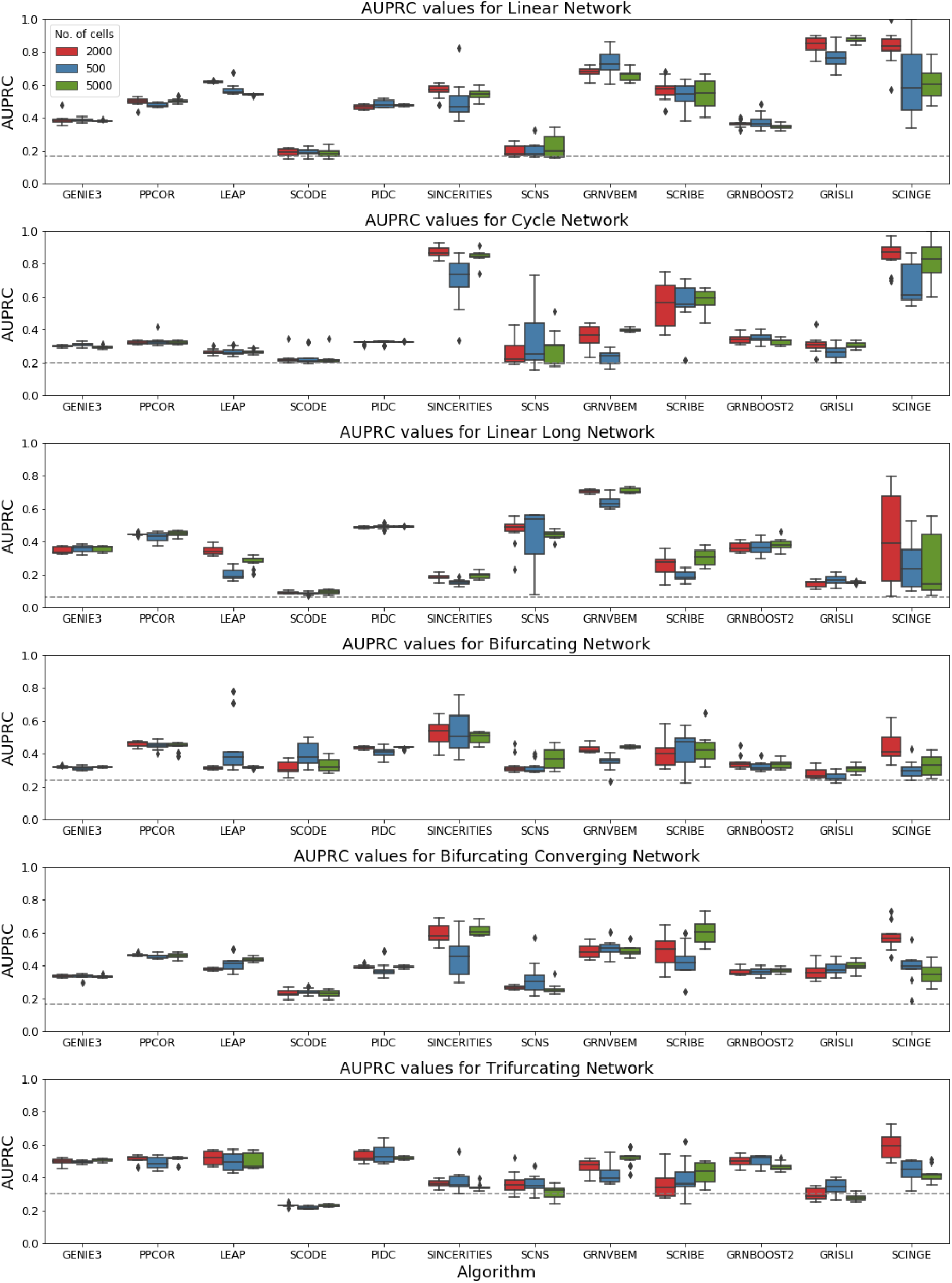
Box plots of AUPRC values. Each row corresponds to one of the six synthetic networks. Each column corresponds to an algorithm. Blue, red, and green box plots corresponds to datasets with 500, 2,000, and 5,000 cells, respectively. The gray dotted line indicates the AUPRC value for a random predictor (network density).

**Figure S3:**
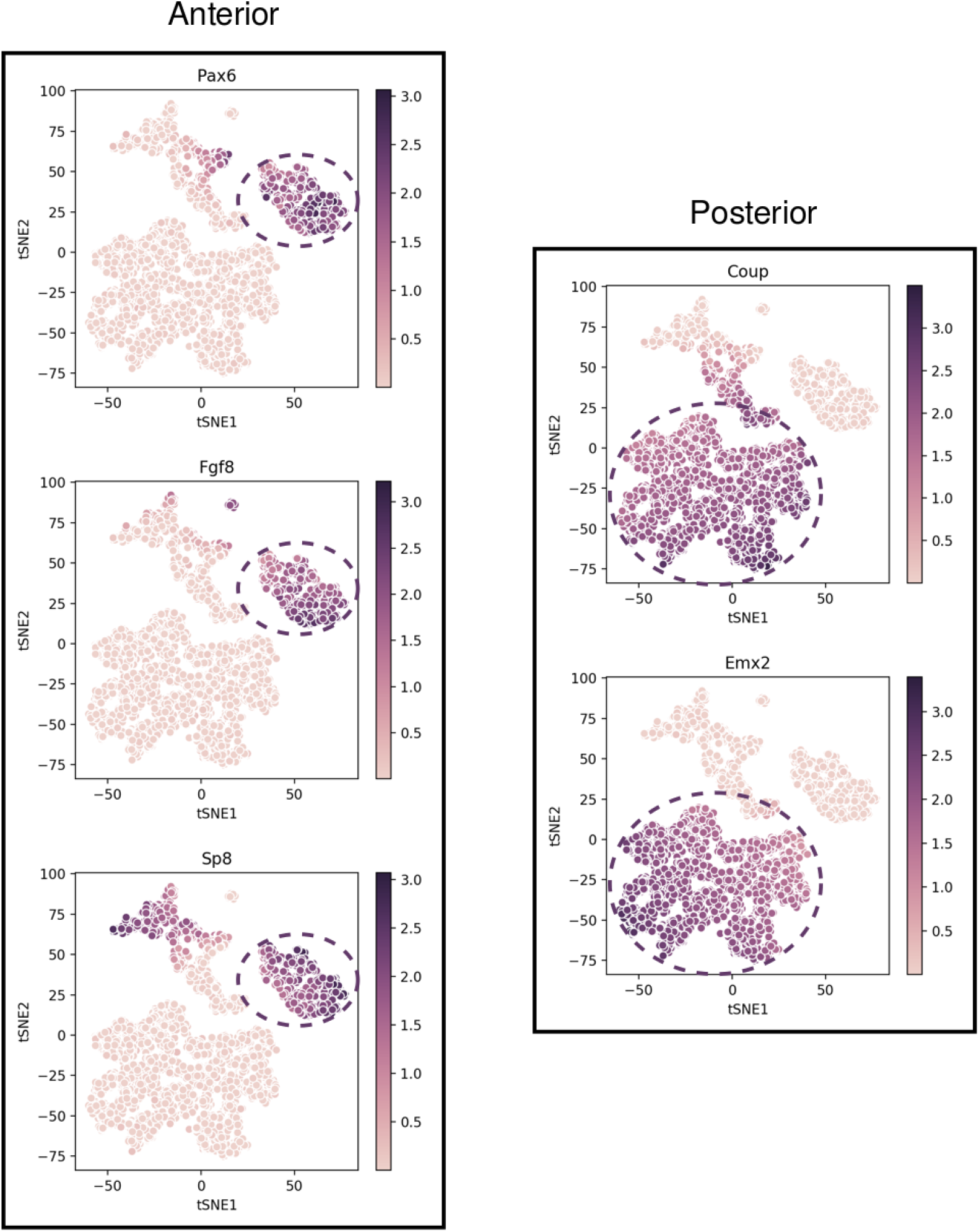
t-SNE visualizations of simulated trajectories from the mCAD model. The bar on the right of each plot displays the mapping between the expression value of a gene and the corresponding color. The model has two steady states corresponding to the anterior and posterior compartments during cortical area development [25]. The anterior compartment is characterized by high transcriptional activity of Fgf8, Pax6 and Sp8, while the posterior compartment is characterized by the high transcriptional activity of Coup-tfi and Emx2. The steady states corresponding to the two compartments appear as two distinct clusters in the t-SNE visualizations.

**Figure S4:**
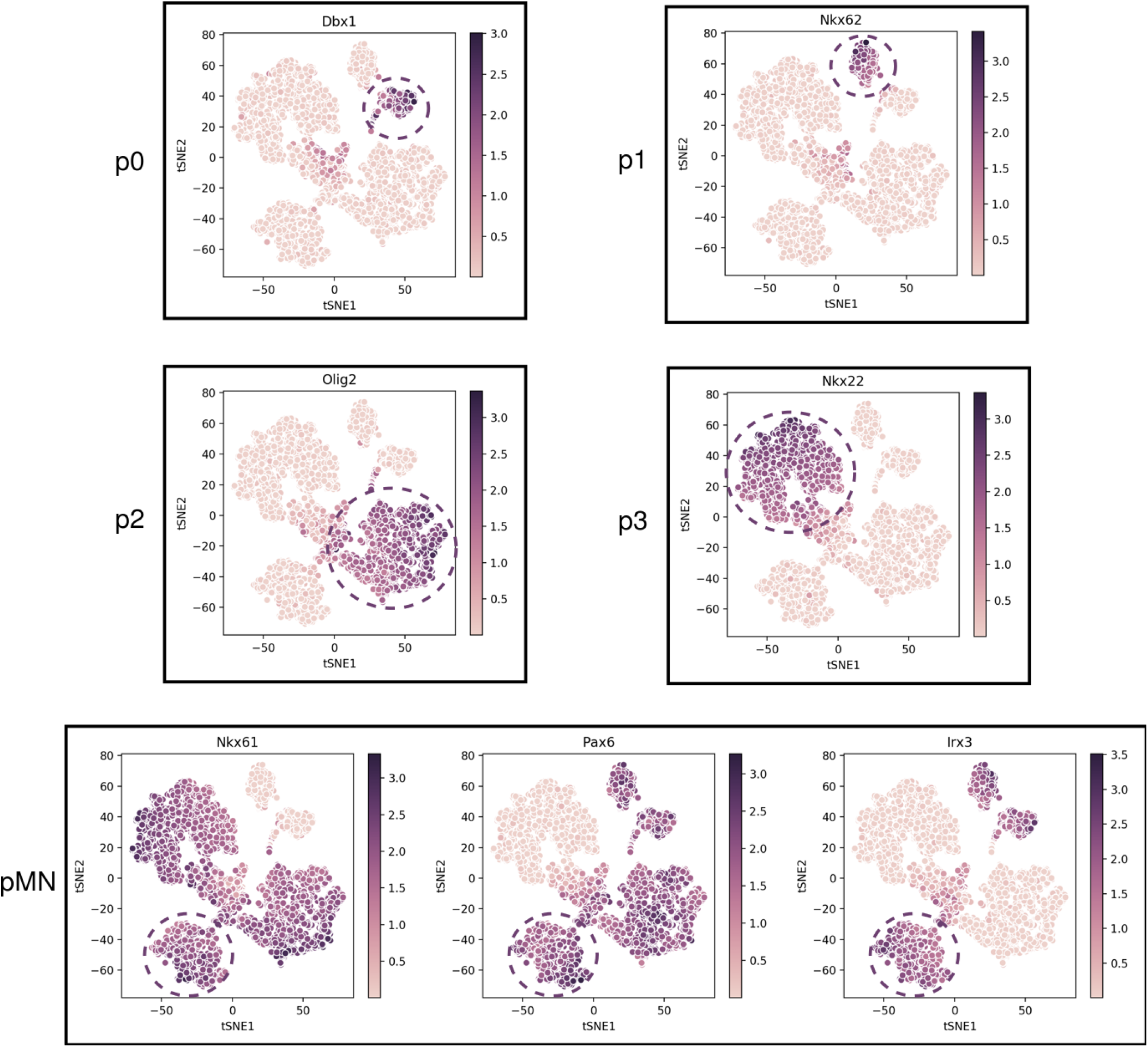
t-SNE visualizations of simulated trajectories from the VSC model. This model exhibits five distinct steady states corresponding to five distinct cell types in the neural tube [26]. Each of these cell types appears as a distinct cluster in the tSNE visualizations shown above. The transcription factors specific to the p0 (Dbx1), p1 (Nkx6.2), p2 (Olig2), and p3 (Nkx2.2) states are shown in the four plots on top. The pMN state is characterized by high activity of Nkx6.1, Pax6 and Irx3, and is shown in the lower plot.

**Figure S5:**
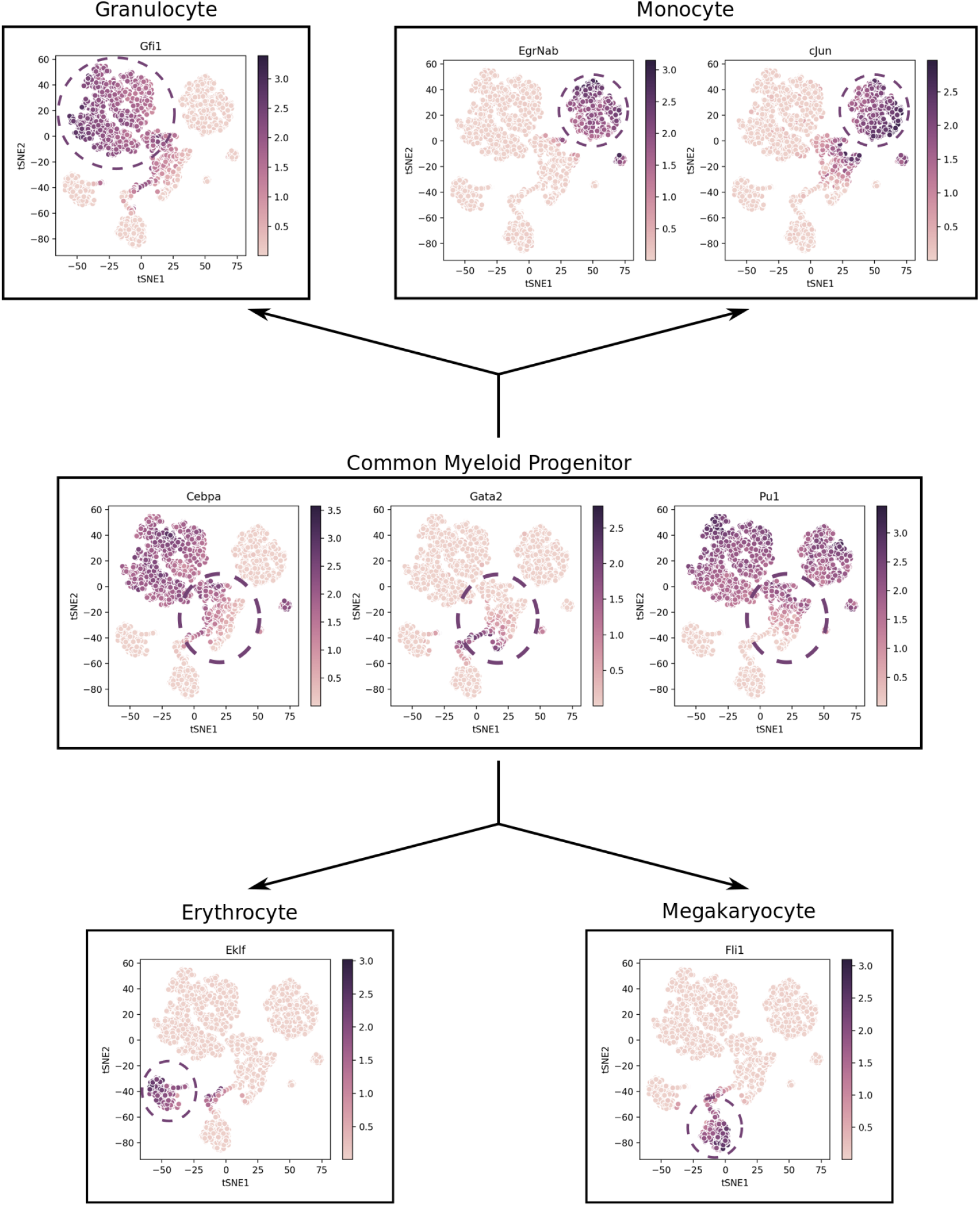
t-SNE visualizations of simulated trajectories from the HSC model. In the initial state, which corresponds to the Common Myeloid Progenitor (CMP), the activities of GATA-2. PU.1 and C/EBP*α* are high [27]. There are two main branches of differentiation. The branch leading to the monocyte-granulocyte progenitors is characterized by the inactivation of GATA-2, followed by the activation of either Gfi1 (granulocyte) or cJun and EgrNab (monocyte) [27]. The clusters in the t-SNE visualization that correspond to these states are shown in the plots on top. Similarly, the branch leading to the megakaryocyte-erythrocyte progenitors is characterized by the inactivation of C/EBPA*α* and PU.1 along with the concomitant activation of GATA1, FOG1 and SCL [27]. The subsequent inactivation of GATA2 along with activation of Fli1 leads to the megakaryocyte state while that wit the activation of EKLF leads to the erythrocyte state [27]. These states are shown below.

**Figure S6:**
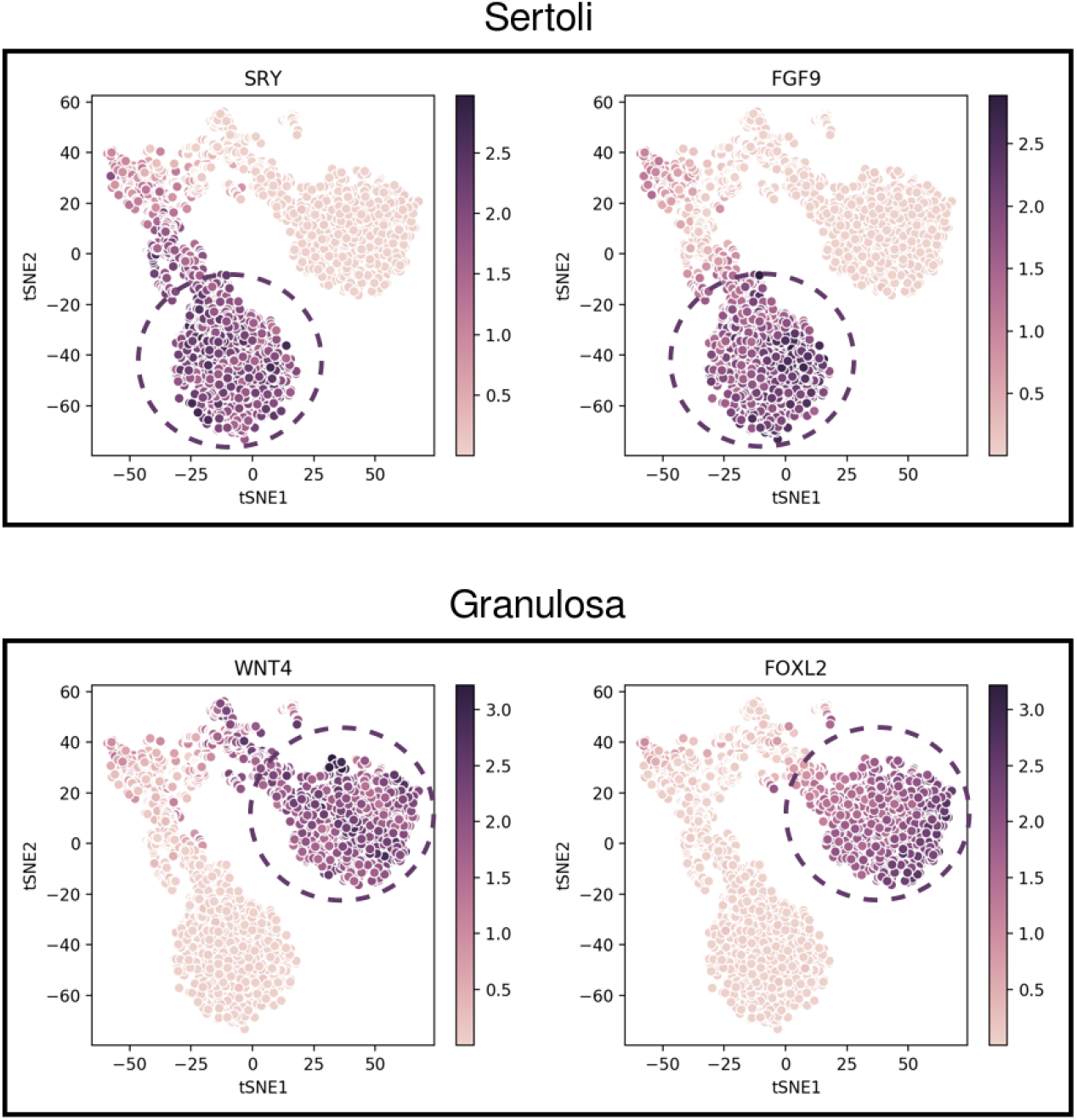
t-SNE visualizations of simulated trajectories from the GSD model. This model has two steady states corresponding to Sertoli cells (male gonads) or Granulosa cells (female gonads) [28]. The clusters on the tSNE plots correspond to these two states. SRY and FGF9 show male-gonad-specific activity while WNT4 and FOXL2 show female gonad-specific activity. Except for GATA4 and WT1mKTS, which are active in both branches, all other transcription factors in the model belong to one of the two clusters shown above.

**Figure S7:**
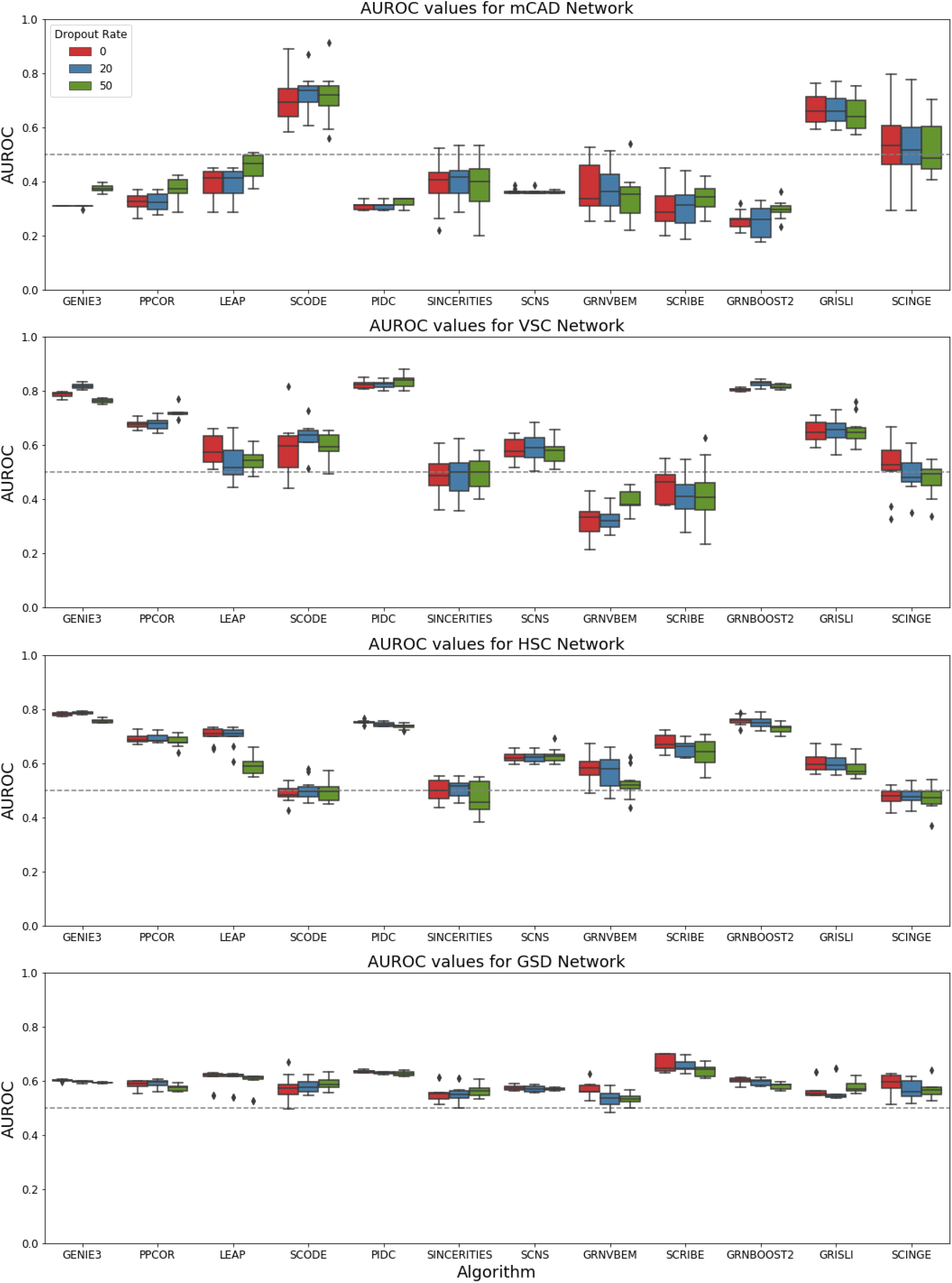
Box plot of AUROC values. Each row corresponds to one of the four synthetic networks. Each column corresponds to an algorithm. Each box plot represents the 10 AUROC values (10 datasets each containing 2,000 cells). Red, blue and green box plots correspond to datasets with no dropouts, a dropout rate of 20%, and a dropout rate of 50%, respectively. The gray dotted line indicates the AUROC value for a random predictor (0.5).

**Figure S8:**
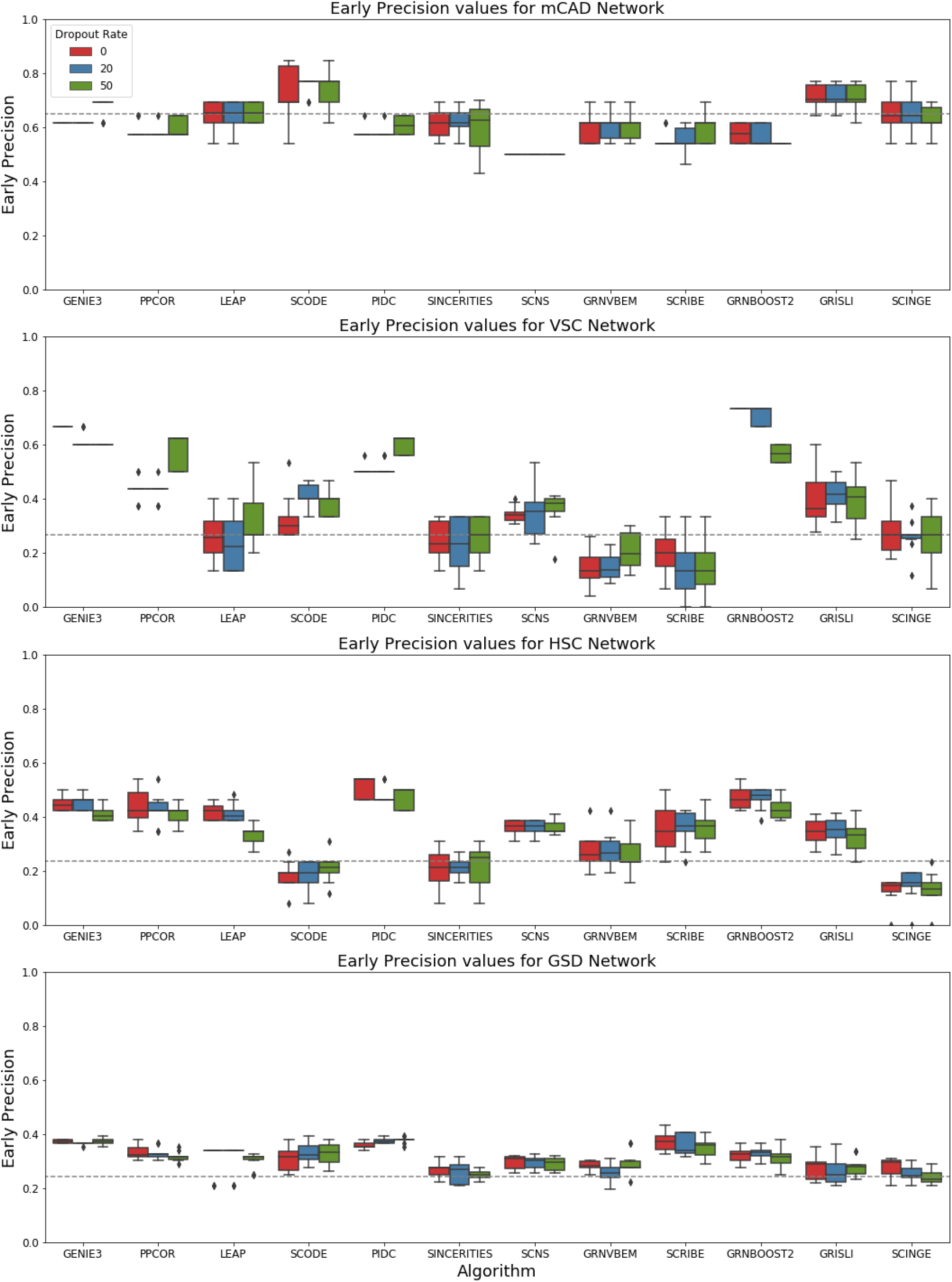
Box plot of early precision values. Each row corresponds to one of the four synthetic networks. Each column corresponds to an algorithm. Each box plot represents the 10 AUROC values (10 datasets each containing 2,000 cells). Red, blue and green box plots correspond to datasets with no dropouts, a dropout rate of 20%, and a dropout rate of 50%, respectively. The gray dotted line indicates the baseline early precision value for a random predictor (network density).

**Figure S9:**
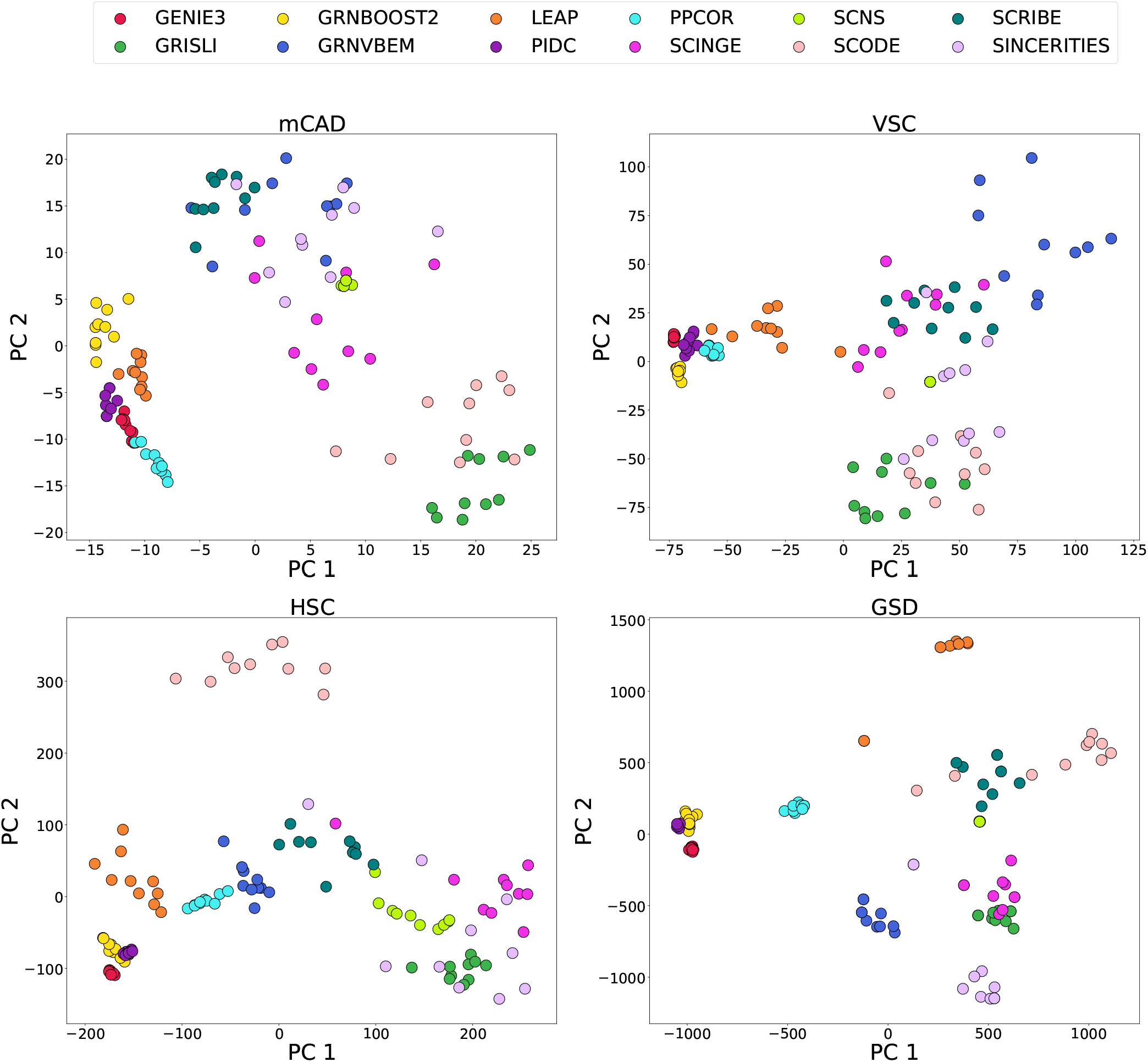
PCA projections of edge rank vectors. Each point corresponds to a vector of edge ranks predicted by an algorithm for a dataset.

**Figure S10:**
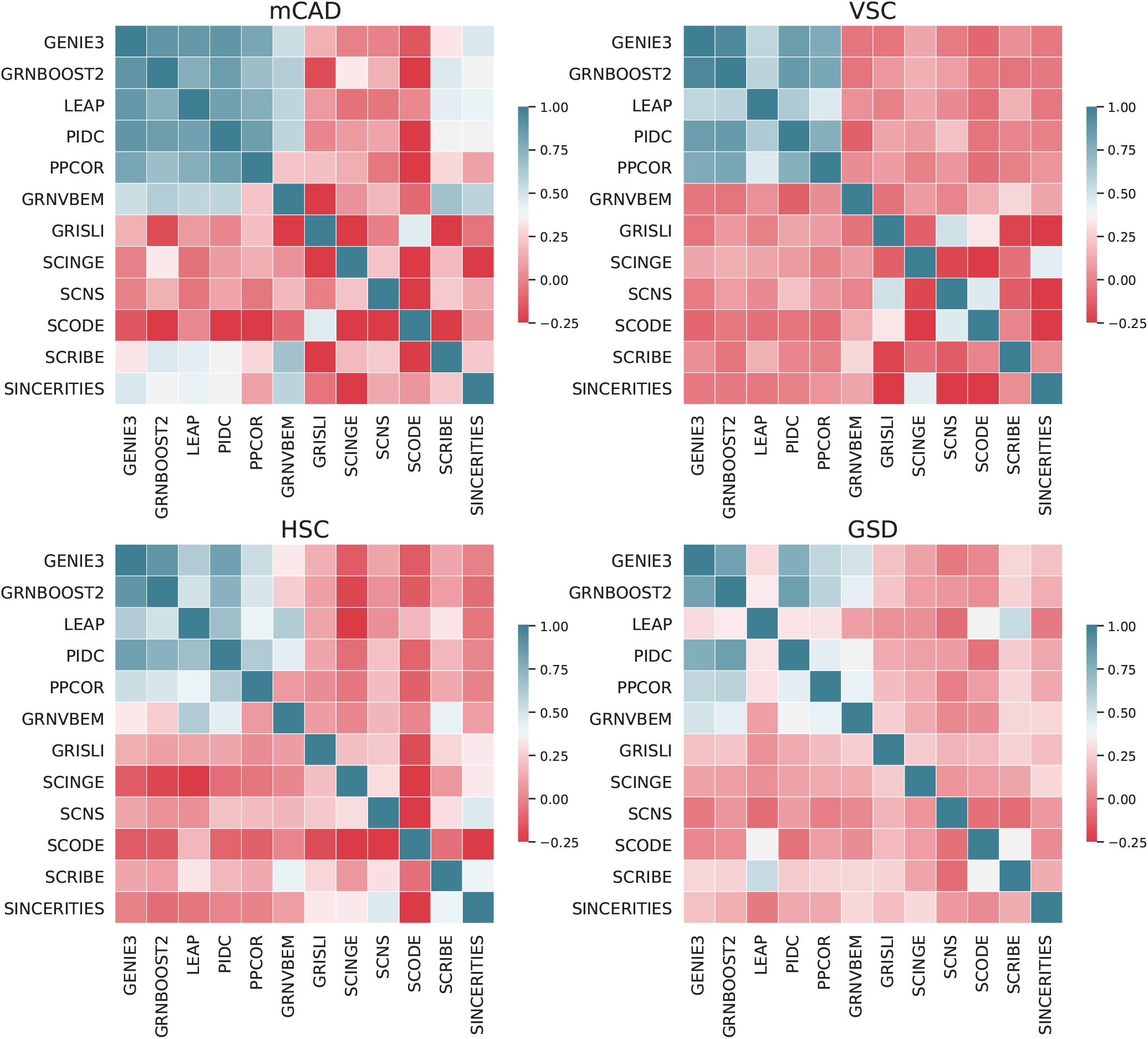
Heatmap of the Spearman correlation matrix for a representative dataset. Each cell represents the correlation value between the edge rank vectors for a pair of algorithms.

**Figure S11:**
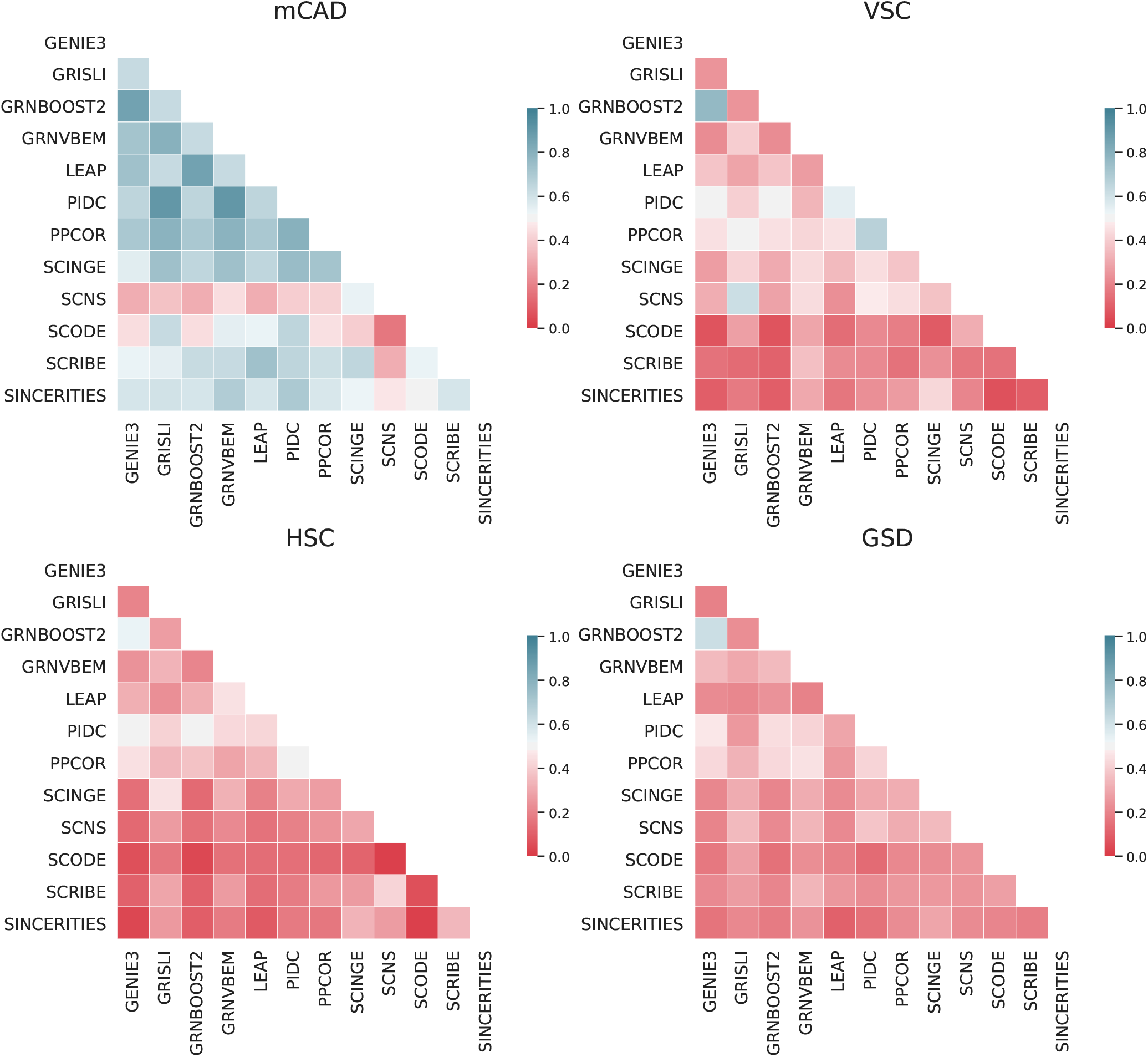
Heatmap of the Jaccard index for a representative dataset. Each cell in the heatmap represents the Jaccard index between the sets of top-*k* edges for a pair of algorithms.

**Figure S12:**
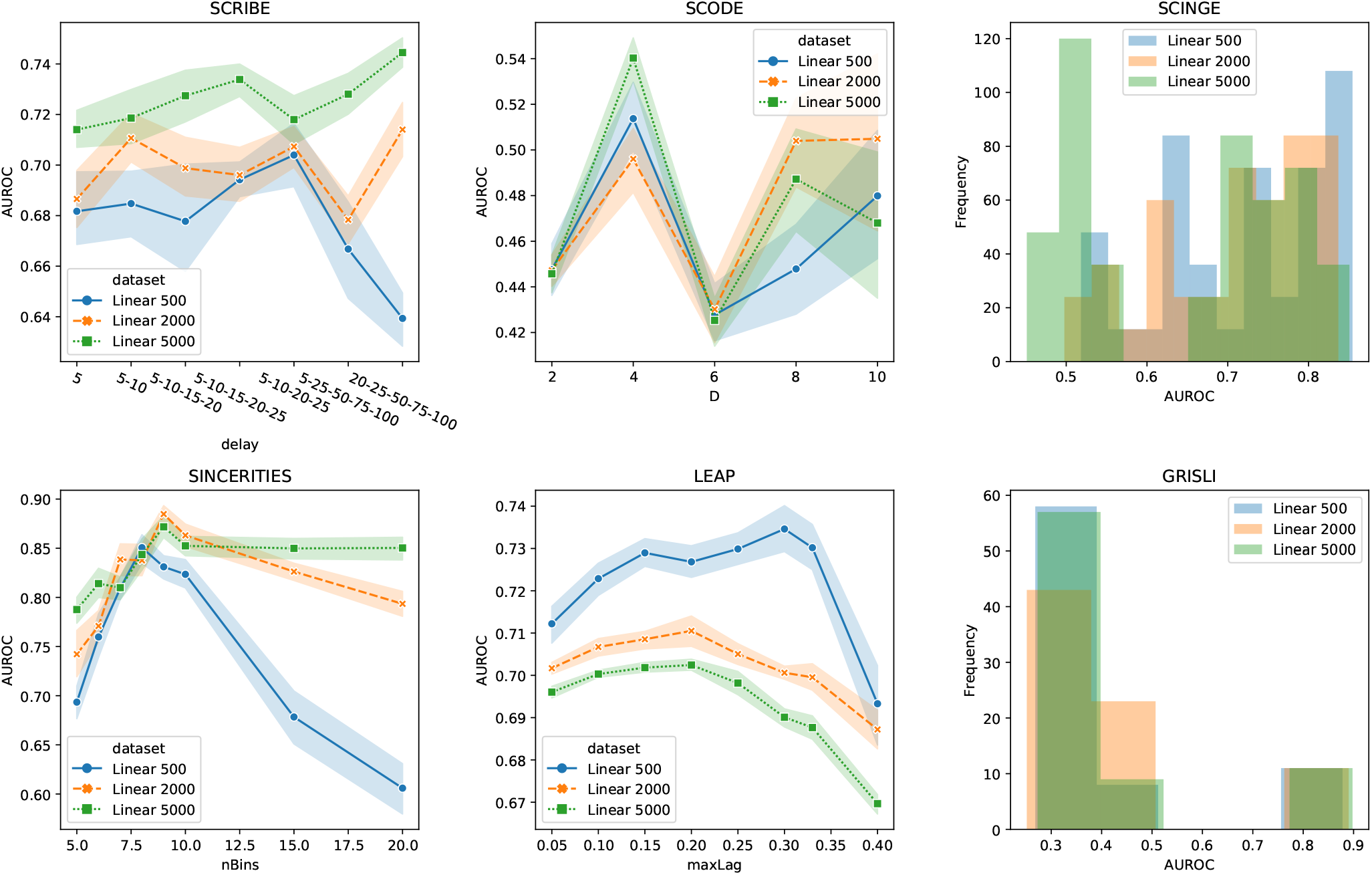
Results of parameter search for each of the six methods with one or more parameters on the datasets (10 each) with 500, 2,000 and 5,000 cells for the Linear synthetic network. For the four methods with one parameter to vary, SCRIBE, SCODE, SINCERITIES, and LEAP, we show the median AUROC as well as the 68% confidence interval of the median, estimated using 1,000 bootstrapping samples of the data. For GRISLI and SCINGE, which have two and seven parameters, respectively, we show histograms of the AUROC over all parameter combinations we tested.

**Figure S13:**
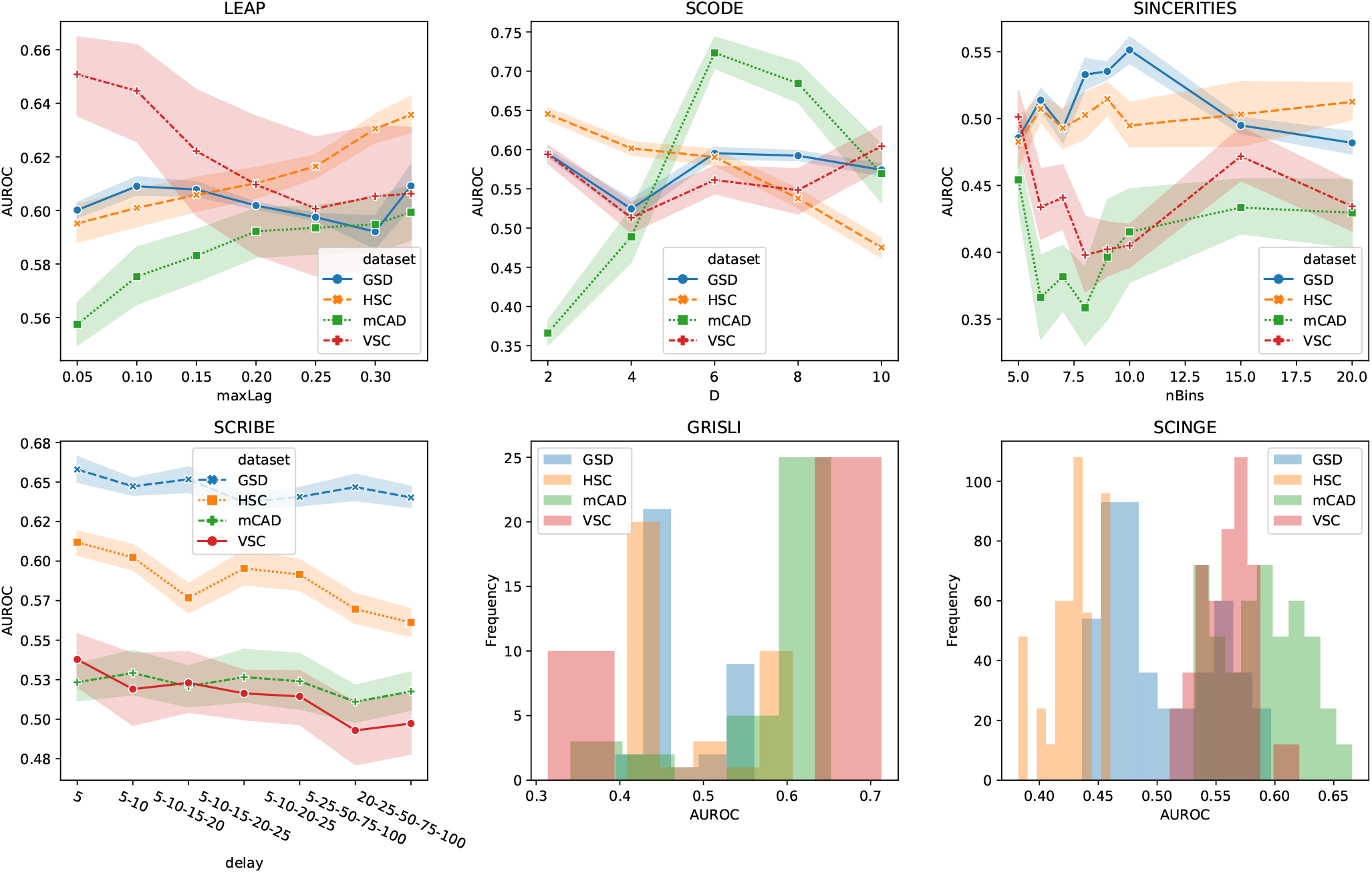
Results of parameter search for each of the six methods with one or more parameters on the 10 datasets with 2000 cells and no dropouts for the four curated Boolean models.

## S5 Supplementary Tables

**Table S2:**
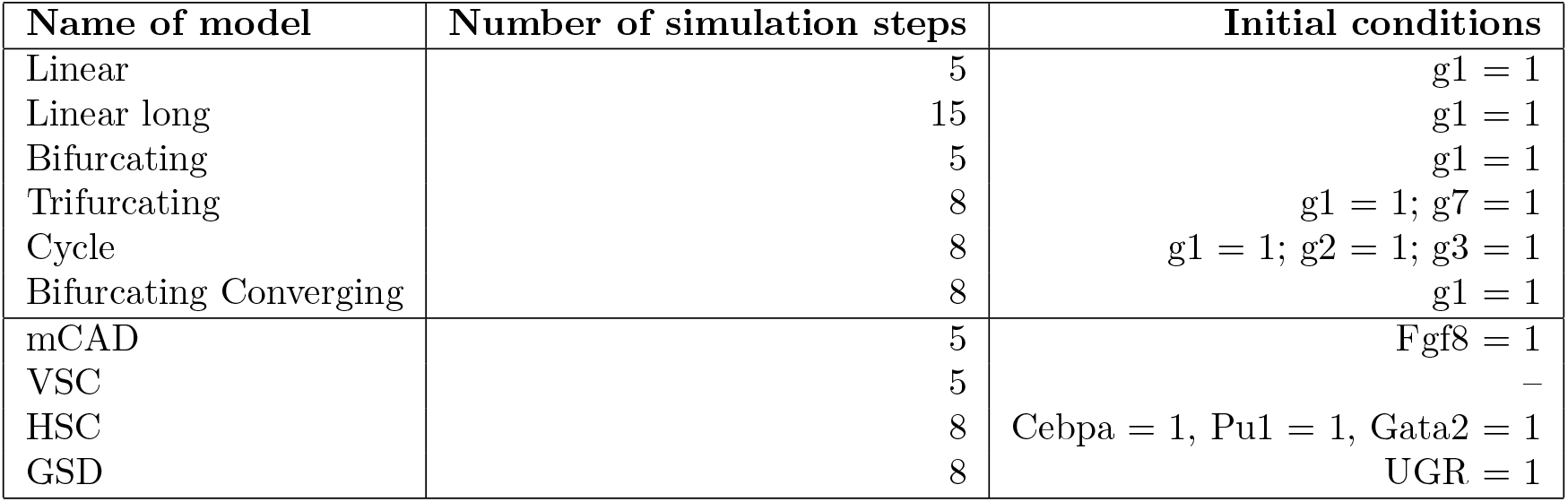
Parameters used to generate synthetic datasets. The final column lists the nodes set to a value of one at the beginning of the simulation. We set all the other nodes to a small value (=0.01) close to zero to avoid numerical errors. For the VCS model, the original publication carried out simulations from all possible states. We carry out VCS simulations by setting the initial conditions of all variables to 1.

| Name of model | Number of simulation steps | Initial conditions |
| --- | --- | --- |
| Linear | 5 | $g1 = 1$ |
| Linear long | 15 | $g1 = 1$ |
| Bifurcating | 5 | $g1 = 1$ |
| Trifurcating | 8 | $g1 = 1; g7 = 1$ |
| Cycle | 8 | $g1 = 1; g2 = 1; g3 = 1$ |
| Bifurcating Converging | 8 | $g1 = 1$ |
| mCAD | 5 | $Fgf8 = 1$ |
| VSC | 5 | – |
| HSC | 8 | $Cebpa = 1, Pu1 = 1, Gata2 = 1$ |
| GSD | 8 | $UGR = 1$ |

